# Stress is basic: ABA alkalinizes both the xylem sap and the cytosol of Arabidopsis vascular bundle sheath cells by inhibiting their P-type H^+^-ATPase and stimulating their V-type H^+^-ATPase

**DOI:** 10.1101/2021.03.24.436813

**Authors:** Tanmayee Torne, Yael Grunwald, Ahan Dalal, Adi Yaaran, Menachem Moshelion, Nava Moran

## Abstract

**BACKGROUND AND HYPOTHESIS:**

• Under water deprivation, in many perennial species, the stress hormone, ABA, appears in the xylem sap in the shoot (including leaf) veins and the xylem sap pH (pH_EXT_) increases. This study aimed to test the hypothesis that ABA is the signal for an altered proton balance of the leaf-vein-enwrapping bundle sheath cells (BSCs).

**METHODS:**

• *Plant Material.* We used a few *Arabidopsis thaliana* (L.) Heynh. genotypes: wildtype (WT) of two accessions, Landsberg *erecta* (Ler) and Columbia (Col), and a few mutants and transformants in these backgrounds.
• *H^+^-Pumps activities.* We monitored ABA effects on the H^+^-pump activities in the BSCs cytosol-delimiting membranes (plasma membrane and tonoplast) by monitoring the cytosol and the xylem pH, and the membrane potential (E_M_), by imaging the fluorescence of pH- and membrane potential (E_M_)-reporting probes: (a) the BSCs’ pH_EXT_ – with the ratiometric fluorescent dye FITC-dextran petiole-fed into detached leaves in unbuffered xylem perfusion solution (XPS), (b) the BSCs’ pH_CYT_ – with the ratiometric dye SNARF1 loaded into BSCs isolated protoplasts, and (c) the BSCs’ E_M_ – with the ratiometric dye di- 8-ANEPPS.

**RESULTS:**

• ABA increased the pH_EXT_; this response was abolished in an *abi1-1* mutant with impaired signaling via a PP2C (ABI1) and in an *aha2-4* mutant with knocked-down AHA2;
• ABA depolarized the WT BSCs;
• ABA increased pH_CYT_ irrespective of AHA2 activity (i.e., whether or not AHA was inhibited by vanadate, or in the *aha2-4* mutant);
• The ABA-induced cytosol alkalinization was abolished in the absence of VHA activity (i.e., when VHA was inhibited by bafilomycin A1, or in the *vha-a2 vha-a3* double mutant with inactive VHA);
• All these results resemble the ABA effect on GCs;
• In contrast to GCs, AHA2 and not AHA1 is the ABA major target in BSCs;
• Blue light (BL) enabled the response of the BSCs’ VHA to ABA;
• The ABA- and BL-signaling pathways acting on both BSCs’ pumps, AHA2 and VHA, are likely to be BSCs autonomous, based on (a) the presence in the BSCs of many genes of the ABA- and BL-signaling pathways and (b) ABA responses (depolarization and pH_CYT_ elevation) demonstrated under BL in isolated protoplasts.

**SIGNIFICANCE STATEMENT:** We reveal here an alkalinizing effect of the plant drought-stress hormone ABA on the pH on both sides of the plasmalemma of the vein-enwrapping bundle sheath cells (BSCs), due to ABA inhibition of the BSCs’ AHA2, the plasmalemma H^+^- ATPase and stimulation of VHA, their vacuolar H^+^-ATPase. Since pH affects the BSCs’ selective regulation of solute and water fluxes into the leaf, these H^+^- pumps may be attractive targets for manipulations aiming to improve plant drought response.

## INTRODUCTION

Bundle sheath cells (BSCs) enwrap the leaf veins in a tight layer which is increasingly recognized as a selective barrier for ions traveling between the leaf veins and the mesophyll (Shapira et al., 2009; Shapira et al., 2013; Wigoda et al., 2017; Niu et al., 2018). This selectivity has been attributed to the membranes of the BSCs, but although the ion transport properties of these cells might impact the plant mineral nutrition and the plant’s susceptibility to various stresses (Wigoda et al., 2014), for example, salt stress or water deficiency stress, they are still severely understudied.

Water deprivation, either gradually in the soil, or by more abrupt changes in the atmospheric VPD, elicits the appearance of the stress hormone ABA in the shoot (Waadt et al., 2014) leading to leaf stomata closure within minutes (Grondin et al., 2015). In Arabidopsis, the source of ABA (as well as its synthesis) has been traced to xylem parenchyma cells enclosed within the BSCs layer (McAdam et al., 2016). Past work from one of our groups (Shatil-Cohen et al., 2011) demonstrated that ABA, fed via a petiole to the xylem of detached Arabidopsis leaves, decreased the leaf hydraulic conductance (K_leaf_) and ABA applied in the bath to isolated BSCs protoplasts, decreased their osmotic water permeability (P_f_). Recently, we reported that the xylem sap pH (pH_EXT_) regulates water access to the leaf across the BSCs, as reflected in the K_leaf_. We showed that xylem sap acidification (by pH buffers; Grunwald et al., 2021) or by blue light which – via PHOT receptors – stimulated the the BSCs’ H^+^-ATPase AHA2 (Grunwald et al., 2020), increased the K_leaf_, while xylem sap alkalinization (by a pH buffer or an absence of blue light) decreased it, and, in parallel, bath alkalinization decreased the P_f_ of isolated BSCs’ protoplasts (Grunwald et al., 2021). Thus, pH_EXT_ affected K_leaf_ by regulating the individual BSCs’ P_f_ (Grunwald et al., 2021; Grunwald et al., 2020), likely through an altered activity of aquaporins in their plasma membrane (Shatil-Cohen et al., 2011; Sade et al., 2014; Sade et al., 2015; Harayama et al., 2019) .

Xylem sap alkalinization within leaf veins occurs in many perennial plants undergoing water deprivation (drought or its mimic), accompanied by the abovementioned increase of ABA in the leaf xylem sap (Wilkinson and Davies, 2008). Similarly, in the dicots guard cells (GCs), water deprivation *raised* the pH_EXT_ (i.e., the apoplastic pH; Geilfus, 2017 and references therein) and this also occurred as the result of direct exogenous ABA application (Felle and Hanstein, 2002). This apoplast alkalinization reflected the ABA-induced cessation of the activity of the plasmalemma (PM) P-type H^+^-ATPase, AHA1 (Hartung et al., 1988). The same treatments – drought or ABA application – led also to the alkalinization of the guard cells cytosol (Irving et al., 1992; Gonugunta et al., 2009; Bak et al., 2013). This was attributed to the ABA-induced stimulation of the activity of the vacuolar V-type H^+^-ATPase, VHA, the underlying details of which are still not completely resolved (Jezek and Blatt, 2017). In Arabidopsis whole etiolated seedlings, and in particular, in the epidermal cells of seedlings roots, applied ABA also alkalinized the apoplast by inhibiting the root PM H^+^-ATPase, but, in contrast to guard cells, it *acidified* the cytosol (Beffagna et al., 1997; Planes et al., 2015).

We chose the best researched guard cell model to guide our exploration of ABA signaling in the BSCs. The ABA signaling cascade leading to the closure of stomata is initiated in the GCs by ABA binding to its intracellular receptor(s) (PYR/PYL), recruiting a clade A protein phosphatase PP2C (e.g., ABI1) to the complex and relieving the inhibition of kinases downstream (Ma et al., 2009; Umezawa et al., 2009). A number of steps downstream, the GCs alkalinize their apoplast (Geilfus, 2017) and their cytosol (Blatt and Armstrong, 1993). In *abi1-1* plants (Koornneef et al., 1984; Meyer et al., 1994), the mutated PP2C protein (*abi1-1*) cannot be recruited to the ABA/PYR/PYL complex upon ABA binding (due to an amino acid substitution G180D, Leung et al., 1997), but still retains its phosphatase activity (Merlot et al., 2001; Imes et al., 2013). The mutation is dominant-negative: it allows the circulation of an active protein phosphatase allowing it to continue its negative regulation of the downstream ABA signaling pathway (Gosti et al., 1999; Cai et al., 2017 and references therein), rendering the guard cells insensitive to ABA (Wu et al., 2003) and eventually preventing the GC’s AHA1 inhibition by ABA.

Here, we test the hypotesis that the water-deficiency-signaling-ABA is what evokes the xylem sap (pH_EXT_) alkalinization and, that in parallel, ABA generates an additional pH signal within the BSCs’ cytosolic milieu (pH_CYT_).

Indeed, we find that in Arabidopsis, exogenous ABA applied to detached leaves via petioles elicits xylem alkalinization, linking these two phenomena in a causative sequence. We demonstrate also an ABA-induced alkalinization of the BSCs cytosol. Furthermore, we single-out the two major proton pumps underlying these phenomena. Finally, we demonstrate that these ABA-induced pH changes are attributable – at least in part – to a BSCs-autonomous ABA signaling cascade, likely to resemble the guard cells’ one.

## RESULTS

### ABA alkalinizes the xylem perfusate in detached leaves

We determined the pH of the xylem perfusate (pH_EXT_) from the fluorescence images of the leaf veins perfused with the ratiometric pH reporter, FITC-D (Materials and methods), as described recently (Grunwald et al., 2021), based on a newly-repeated calibration curve (Suppl. Fig. S1). A non-bufferred xylem perfusion solution (XPS, Solutions, Materials and methods), imbibed via the petiole for about 1.5-2 h into detached mature leaves of WT (Ler) Arabidopsis, attained a pH of roughly 6.7 (Fig. 1A, Suppl. Fig. S1). When 10 M ABA was included in the XPS, pH_EXT_ increased by about 0.4 pH units (from pH 6.7 to 7.1; Fig. 1A), suggesting that ABA inhibited proton extrusion from the BSCs into the xylem by inhibiting the BSCs AHA.

**FIGURE 1.**
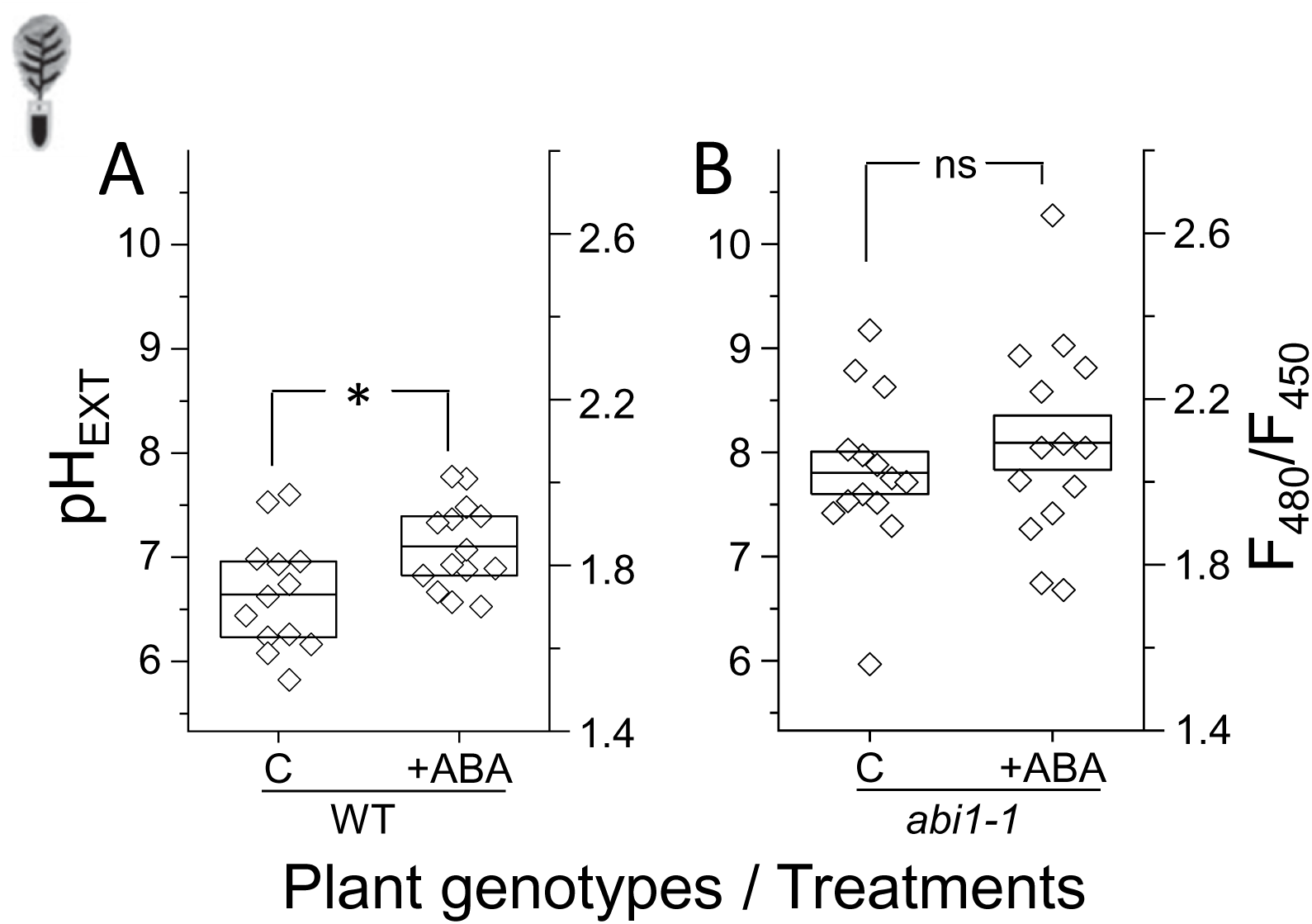
A protein phosphatase 2C (PP2C, aka ABI1) mediates the ABA-induced alkalinization of the xylem perfusate in the minor veins of Arabidopsis detached leaves. **A:** The effect of ABA (10 μM) on the xylem sap pH (pH_EXT_) in WT *Arabidopsis thaliana,* accession Landsberg *erecta* (Ler). The data are from at least three independent experiments. Asterisk: significant difference (at P=0 0.02048, by Student’s 2-tailed t-test, equal variance). Other details as in Suppl. Fig. S3A. **B:** ABA effect on the pH_EXT_ of the *abi1-1* (*ABA insensitive*, Ler) mutant’s leaves. ns: non-significant difference. Note the absence of ABA effect in the absence of the phosphatase ABI1 activity. Inset: a detached leaf schematic.

### Does ABA signaling in the BSCs occur via the protein phosphatase ABI1?

To resolve this, we compared the ABA response of the XPS in the WT (Ler) plants to that in *abi1-1* (Ler) plants, i.e., plants harboring a dominant-negative mutation in the gene of of the clade A protein-phosphatase 2C (PP2C, aka ABI1; Materials and methods), isolated initially by (Koornneef et al., 1984) and further characterized by (Leung et al., 1997). In the ABA-treated *abi1-1* (Ler) mutants leaves, the XPS pH remained unaltered (pH_EXT_ around 7.8-8.1; Fig. 1B). Similarly, ABA induced XPS alkalinization in the WT (Col) by about 0.8 units (pH_EXT_ change from 6.5 to approx. 7.3; Suppl. Fig. S2A), while failing to affect pH_EXT_ (it remained unchanged around 6.6-6.8; Suppl. Fig. S2B) in the leaves of *SCR: abi1-1* (Col) plants, generated by introducing the *abi1-1* gene into WT (Col) under the BSCs-directing Scarecrow (SCR) promoter (Suppl. Figs. S3A, S3B; Materials and methods). These results suggested the necessity of an intact ABI1 (in the BSCs, and / or in their vicinity, see Discussion) for ABA alkalinization of the XPS. Interestingly, the stomata in the epidermal peels of the *SCR:abi1-1* plants remained fully responsive to ABA (Suppl. Fig. S3C).

### Is AHA2 involved in the ABA-induced XPS alkalinization in detached leaves?

Earlier, we established the BSCs’ AHA2 as a major xylem sap acidifier (Grunwald et al., 2021; Grunwald et al., 2020). To resolve whether AHA2 is involved also in the ABA-induced XPS alkalinization, we compared the ABA responses in detached mature leaves of WT (Col) Arabidopsis to those in the *aha2-4* mutant plants. *aha2-4* is a knockdown mutant of AHA2 (Haruta et al 2010, 2012). Similar to the previous experiments (Fig. 1A, Suppl. Fig. S2A), ABA alkalinized the XPS in the WT (Col) leaves, by 0.4 pH units (from 5.8 to 6.2, Fig. 2A). In contrast, ABA did not affect significantly the pH_EXT_ in the *aha2-4* leaves (it remained in the range of 6.0 to 5.7; Fig. 2B), suggesting that in the absence of AHA2 activity, ABA did not affect proton extrusion via the plasma membrane. However, in *aha2-4* complemented with *SCR:AHA2*, restoring the missing AHA2 activity exclusively in the BSCs (Grunwald et al., 2021), ABA treatment increased the pH_EXT_ by 0.8 units, doubly that in WT (from 5.6 to 6.4; Fig. 2C), indicating that in the BSCs, AHA2 is a target of ABA-induced inhibition.

**FIGURE 2.**
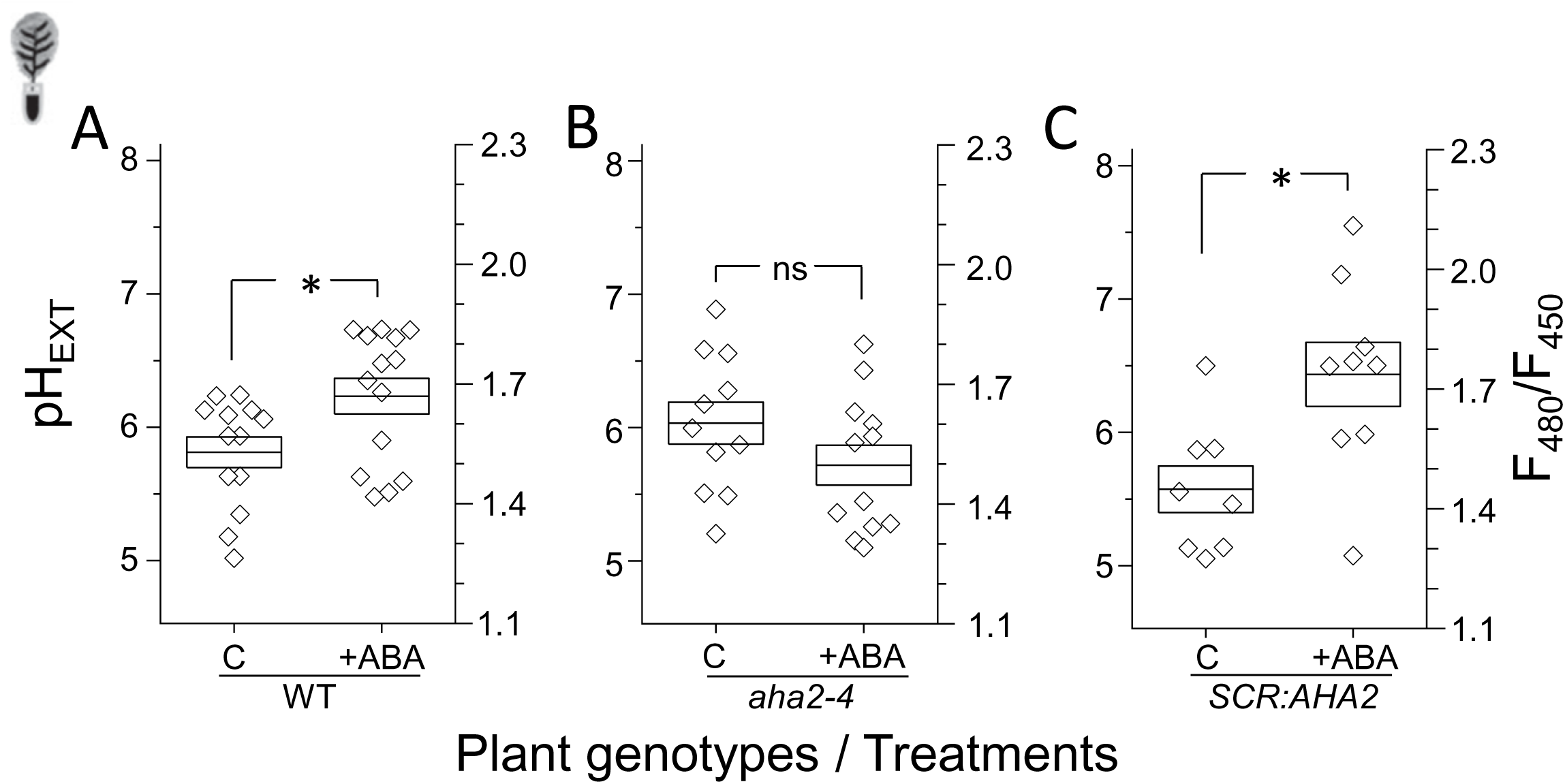
ABA alkalinization of the xylem sap in the minor veins of detached leaves results from the cessation of the activity of the BSCs’ AHA2. **T**he effect of ABA on the pH of XPS (pH_EXT_) in the leaves of Arabidopsis (Col) with and without AHA2 activity, derived from F-ratio images (Materials and methods, Grunwald et al., 2020). **A.** WT: wild type Arabidopsis (Col). Asterisk: significant difference (at P=0.02596, by 2-tailed t-test, equal variance). The data are from at least three independent experiments. B. *aha2-4* (Col), a line of a knockdown mutant of the AHA2 (Haruta et al., 2010, 2012). ns: a non-significant difference. Other details as in A. C. *SCR:AHA2*: *aha2-4* complemented with *SCR:AHA2* exclusively in their BSCs (Grunwald et al., 2020). Asterisk: a significant difference (at P=0.01238). Other details as in A. Inset: a detached leaf schematic. Note the absence of ABA effect in the absence of AHA2 activity.

### The effect of ABA on the BSCs membrane potential, E_M_

In addition to largely driving the apoplast pH (pH_EXT_), the electrogenic proton extrusion of AHA2 hyperpolarizes the plasma membrane (reviewed by Spanswick, 1981; Grunwald et al., 2020) and participates in the cytosolic pH homeostasis (Sze and Chanroj, 2018 and references therein). Therefore, the predicted outcome of ABA-induced cessation of proton extrusion at the single-cell level would be the acidification of the BSCs’ cytosol and membrane depolarization, both due to an impaired balance of proton fluxes across the plasma membrane resulting in a net proton influx into the cytosol (as reviewed by Sze and Chanroj, 2018). To examine these predictions, we resorted to assays at a single cell level, which allowed monitoring of very rapid ABA responses, within 5 to 15 minutes.

To test the prediction that ABA depolarizes the BSCs, we applied 3 ☐M ABA to BSCs protoplasts isolated from WT plants bathing in a low-K^+^ solution (Materials and methods) and monitored their membrane potential, E_M_, by the fluorescence of the ratiometric, dual-excitation dye, di-8-ANEPPS (Wigoda et al., 2017; Grunwald et al., 2020; Materials and Methods). As expected, ABA depolarized the BSCs of WT plants by about 100 mV (from about -300 mV to about -200 mV (Fig. 3, calibrated as in Grunwald et al., 2020)

**FIGURE 3.**
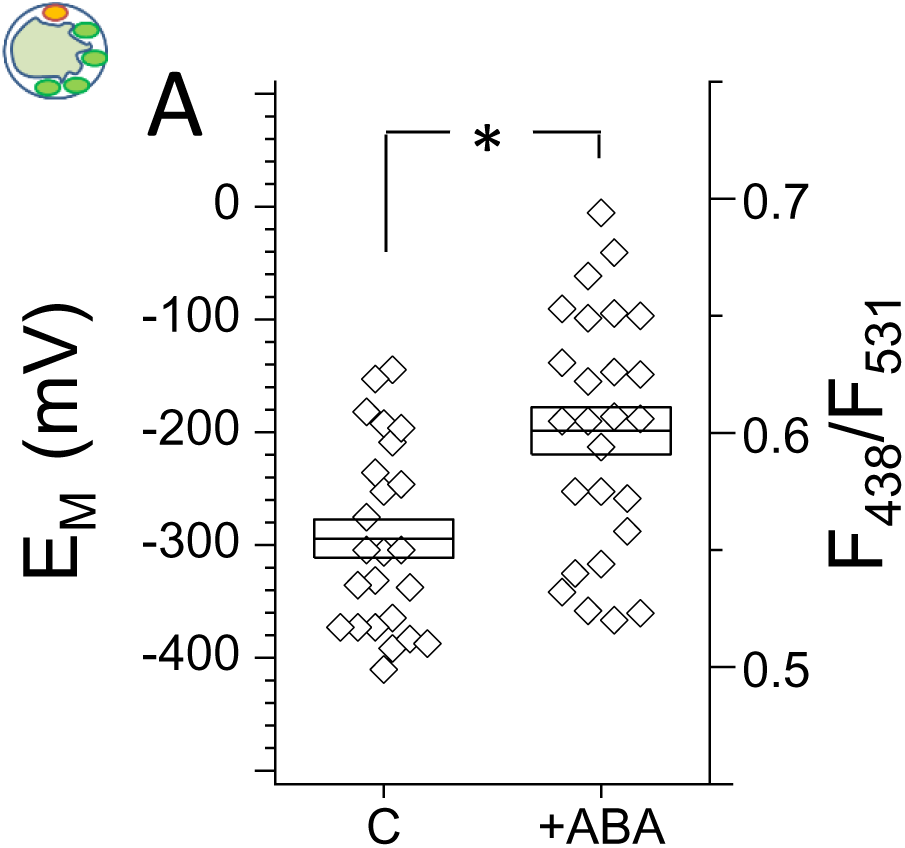
ABA depolarizes the WT (Col) BSCs. Membrane potential (E_M_) of BSCs protoplasts (Col) determined using the fluorescent dual-excitation (Ex_1_: 438, Ex_2_: 531, Em: 550 nm) ratiometric dye, di-8- ANEPPS (30 μM, Materials and methods). BSCs protoplasts, labeled with *SCR:GFP* (to enable their GFP-fluorescence-based selection under the microscope, Materials and methods) were bathed in a low K^+^ solution (5 K, pH 5.6, see Solutions) without (C) or with (+) 3 μM ABA. Shown are the E_M_ values of individual BSCs (symbols, biological repeats) and their means ± SE (midline and box), from at least three independent experiments. The E_M_ values on the ordinate are based on an *in-situ* calibration curve obtained using patch-clamp (Materials and methods, see also Grunwald et al., 2020). Asterisk: a significant difference (at P=0.00095, two-tailed t-test, equal variances). Inset: a protoplast schematic.

### Is AHA2 involved in the ABA-induced response of the BSCs’ pH_CYT_?

To test the prediction that AHA2 inhibition by ABA acidifies the cytosol, we monitored the cytosolic pH (pH_CYT_) using the ratiometric, dual-emission fluorescent pH-sensitive dye, SNARF1 (Materials and methods, Suppl. Fig. S4). Contrary to this simple prediction, 3 ☐M ABA added in the bath with isolated WT (Col) BSCs protoplasts, did not acidify, but rather, alkalinized the cytosol in WT by 0.9 pH units within 0-15 minutes (from 7.7 to 8.6, Fig. 4A), suggesting that the BSCs’ cytosol alkalinization does not depend on the level of their AHA activity. However, in difference to WT and in accord with the above prediction, the “control” *aha2-4* mutant’s pH_CYT_ was indeed lower than the WT’s pH_CYT,_ by 1.4 pH units (the mutant’s 6.3 vs WT’s 7.7; Fig. 4A). Moreover, treating the *aha2-4* mutant BSCs with ABA replicated the effect found in WT, causing an even more pronounced alkalinization, by 1.2 pH units (roughly from 6.3 to 7.5, Fig. 4A), definitively ruling out the involvement of AHA2 in the ABA-induced BSCs’ cytosol alkalinization.

**FIGURE 4.**
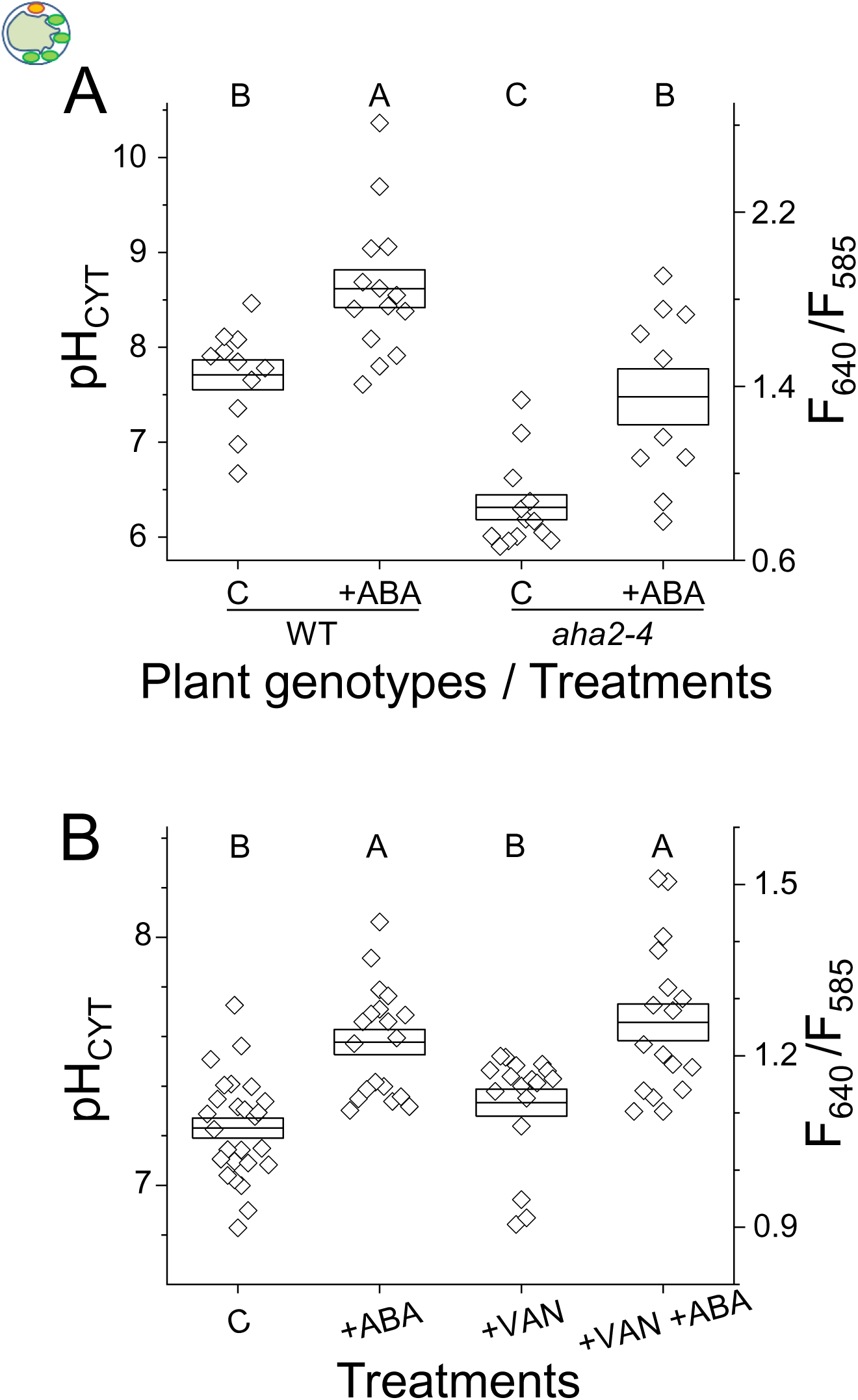
AHA2 is not involved in BSCs alkalinization by ABA. Cytosol pH (pH_CYT_) of BSCs protoplasts determined using SNARF1 (Suppl. Fig. S4; Materials and methods). **A.** The effect of *AHA2* knockdown on ABA response. WT (Col) BSCs and *aha2-4* (Col) mutant plants BSCs were bathed in a low K^+^ solution (see Solutions) without any additive (C) or with (+) 3 μM ABA (Materials and methods). Shown are the individual BSCs’ pH_CYT_ values (symbols, biological repeats) and their means ± SE (midline and box), from at least three independent experiments. The pH_CYT_ values were determined from the F-ratio (F_640_ /F_585_) values, based on the *in-situ* calibration curve and conversion relationship of Suppl. Fig. S4C. Different letters indicate statistically different means (at P<0.05, ANOVA, Tukey HSD test). Inset: a protoplast schematic. **B.** The effect of AHA inhibitor, vanadate, on the ABA response in WT (Col) BSC protoplasts. The protoplasts, bathed as above, were treated without anything (C), or with (+) 3 μM ABA ± 1 mM vanadate (VAN). The basis for conversion of F-ratio to pH_CYT_ was Suppl. Fig. S4B. Other details are as in A. Note that AHA2 activity is not required for the cytosol alkalinization by ABA.

To narrow down the candidacy of other proton pumps potentially responsible for this fast ABA-induced cytosol alkalinization, we pre-incubated the WT (Col) BSCs protoplasts with 1 mM vanadate (a known general P-type ATPases inhibitor; reviewed by Morsomme and Boutry, 2000; Palmgren, 2001 ; Materials and methods). Surprizingly, when we applied vanadate by itself to WT (Col) BSCs, to mimic the *aha2-4* mutation, it did not affect the pH_CYT_ relative to control (C) BSCs (i.e., pH_CYT_ remained in the range 7.2-7.3; Fig. 4B). This contrasted sharply with the pH_CYT_-lowering effect of knocking down AHA2 (which lowered the untreated (C) *aha2-4*’s pH_CYT_ by 1.4 pH units vs. the control (C) WT’s pH_CYT_; Fig. 4A).

Vanadate did not prevent, however, the ABA-induced increase of the BSCs pH_CYT_; with ABA added on top of vanadate in the bath (Materials and methods), pH_CYT_ increased by approximately 0.4 pH units relative to vanadate alone (roughly, from 7.3 to 7.7, Fig. 4B). This ruled out the involvement of any vanadate-susceptible P-type H^+^-ATPases (including the AHA2 (Morsomme and Boutry, 2000; Palmgren, 2001) in the ABA-induced BSCs’ cytosol alkalinization, suggesting, instead, a vanadate-*in*sensitive proton-extruding mechanism, such as, for example, the vacuolar H^+^-ATPase, VHA or the vacuolar pyrophosphatase (PP-ase), APV1 (Krebs et al., 2010).

### Is VHA involved in the ABA-induced cytosol alkalinization?

To explore directly the involvement of VHA in the ABA-induced cytosol alkalinization, we used the VHA-specific inhibitor, bafilomycin A1 (Muroi et al., 1994; Klychnikov et al., 2007; Yano et al., 2015). Curiously, 100 ☐M bafilomycin A1 added to the bath by itself (Materials and methods), elevated the pH_CYT_ within minutes, roughly by 0.4 pH units relative to the control (from about 7.2 to 7.6, Fig. 5A), not much different from the WT’s response to ABA (from 7.2 to 7.5, Fig. 5A). However, when bafilomycin A1 (the VHA inhibitor) was added in the bath together with 3 M ABA (the AHA2 inhibitor), pH_CYT_ rise was abolished (its value remained approx. 7.2, Fig. 5A), suggesting a compensatory action of an ABA- inhibitable H^+^-extrusion mechanism underlying the bafilomycin A1-induced cytosol alkalinization (see Discussion).

**FIGURE 5.**
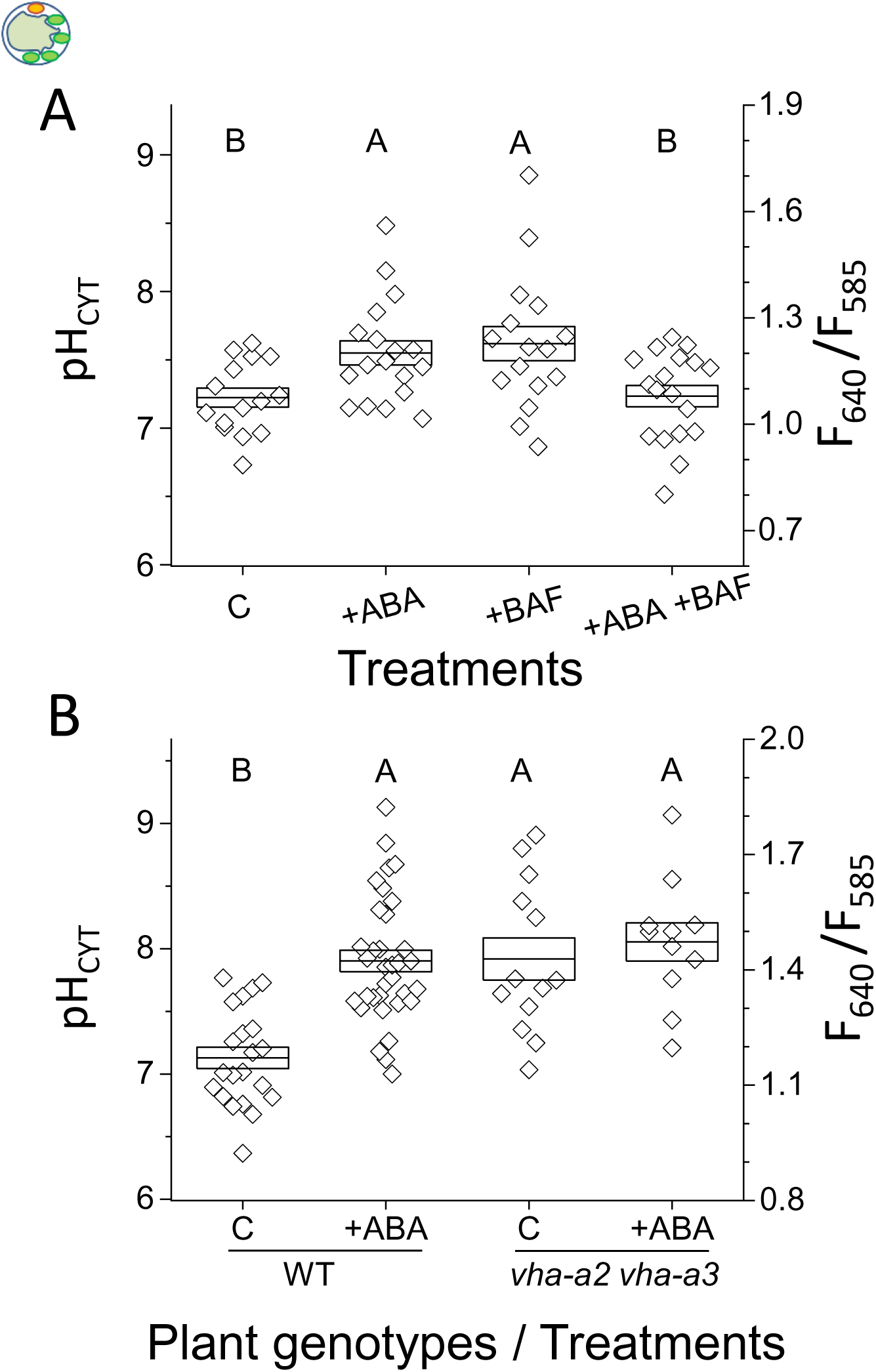
ABA-induced cytosol alkalinization requires the activity of VHA. Cytosol pH (pH_CYT_) of BSCs protoplasts determined using SNARF1 as in Fig. 4. **A.** The effect the specific inhibitor of the vacuolar H^+^-ATPase (VHA), bafilomycin A1 (BAF), on ABA response. The BSCs protoplasts of WT (Col) were bathed in a low-K^+^ solution (see Solutions) without any additive (C) or with (+) 3 μM ABA ± 0.1 μM bafilomycin A1 (BAF; Materials and methods). Different letters indicate significantly different values (at P<0.05, ANOVA, Tukey HSD test). Note the ABA- or BAF- induced pH_CYT_ elevation relative to C, as opposed to their in-effectivity when applied simultaneously. Inset: a protoplast schematic. **B.** The effect of double VHA (Col) mutation *vha-a2 vha-a3*, on the ABA pH_CYT_ response. Note the lack of ABA effect on the pH_CYT_ of the mutant devoid of VHA activity.

In a complementary experiment, we assayed the effect of ABA on the double mutant of the two tonoplast-localized isoforms of the A subunit of VHA (*vha-a2 vha-a3* (Col); Krebs et al., 2010). The “control” pH_CYT_ of the BSCs in this VHA- activity-devoid mutant was more alkaline than in WT (Col), by about 0.7 pH units (roughly, 7.9 vs. 7.2, respectively; Fig. 5B). This may be attributed to the life-long disruption of homeostasis of the vacuolar pH, pH_VAC_ (see Discussion). ABA treatment of the double-mutant did not affect its pH_CYT_ (with pH_CYT_ remaining roughly at 7.9-8.1, Fig. 5B), suggesting that the fast, ABA-induced elevation of pH_CYT_ in the WT requires an active VHA (see Discussion).

### Is light necessary for the BSCs’ VHA activation by ABA?

We learned recently that the BSCs’ AHA2 is activated by light, and, in particular, by blue light added on top of red light (BL+RL) (Grunwald et al., 2020). Further, blue light increased 2-3-fold the enzymatic (hydrolytic) bafilomycin A1-sensitive activity of V-ATPase assayed in tonoplast vesicles of etiolated barley (Klychnikov et al., 2007). Therefore, all our experiments described so far were performed on BL-illuminated BSCs (Materials and methods). To test whether the BSCs’ VHA activity depended similarly to AHA2 activity on light, and, in particular, whether it required substantial illumination to be activated by ABA, we examined the effect of ABA in darkness (≤0.025 ☐mol s^-1^ m^-2^; Materials and methods). In these conditions, ABA did not alkalinize the BSCs’ pH_CYT_ (which remained around 7.6, Fig. 6A), suggesting that ABA did not activate their VHA in the dark.

**FIGURE 6.**
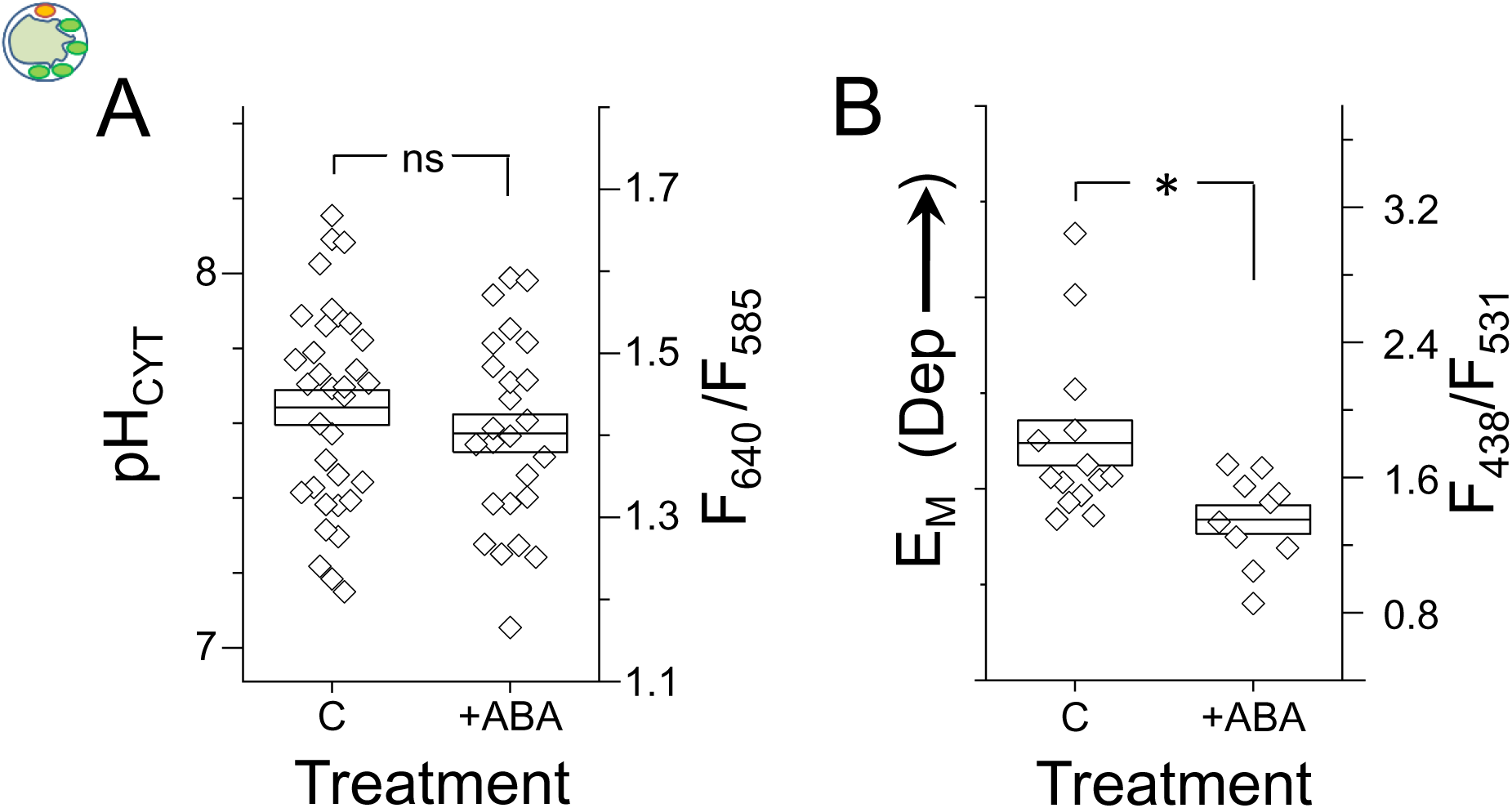
In dim light, ABA does not alkalinize the BSCs’ cytosol nor does it *<u>de</u>*polarize the BSCs. WT (Ler) BSCs protoplasts were bathed in a low K^+^ solution (see Solutions) without the BL+RL illumination (Materials and methods). **A.** The BSCs’ pH_CYT_ without (C) or with 3 μM ABA was determined using SNARF1. Other details as in Fig. 4B. Inset: a ‘protoplast in the dark’ schematic. **B.** The BSCs’ membrane potential (E_M_) was determined using di-8-ANEPPS as in Fig. 3, except for the E_M_ calibration (Materials and methods). Arrow indicates depolarization (Dep). Asterisk: a significant difference (at P= 0.01587, by 2 tailed t-test, equal variances). Other details as in Fig. 3. Note the ABA-induced *<u>hyper</u>*polarization.

**FIGURE 7.**
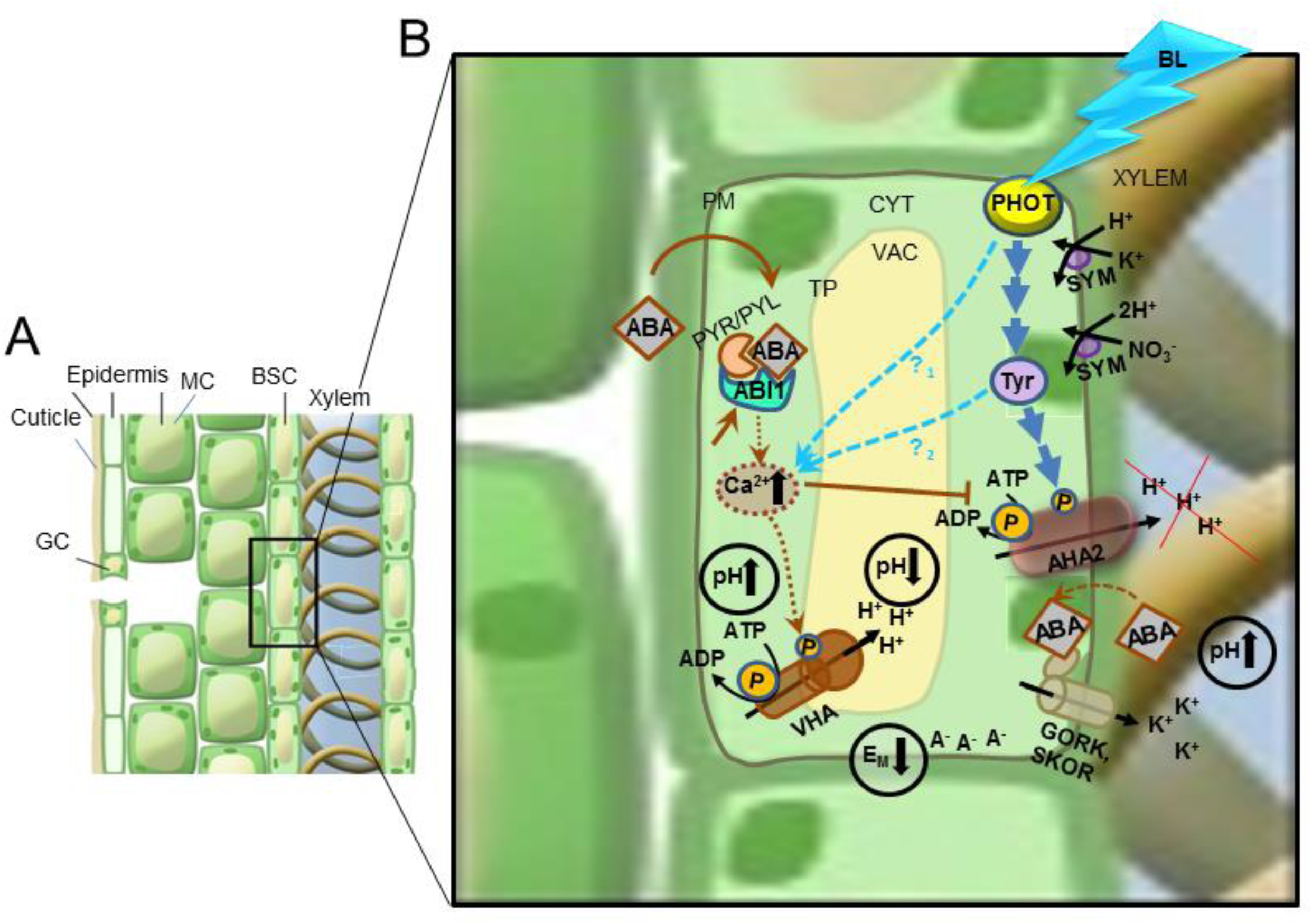
A proposed BSCs-autonomous ABA and BL signal transduction pathways. An artist’s rendering of changes in an ionic balance between the BSCs and the xylem, due to a simultaneous regulation of two H^+^-ATPases, the plasma membrane (PM) AHA2 and the tonoplast (TP) VHA, and the K^+^ -efflux channels, GORK and SKOR. **A.** GC, a stomata guard cell; MC, mesophyll cell. **B.** A BSC in an expanded view. *Dark blue arrows:* the Blue-Light (BL) signalling pathway activating the AHA2, from BL perception by the phototropin receptors (PHOT), through an intermediate tyrphostin (Tyr)-sensitive tyrosine phosphorylation, resulting in H^+^ extrusion and xylem sap acidification (Grunwald et al., 2020; Grunwald et al., 2021). *Light blue arrows:* the BL signalling pathway ‘primes’ the V-type H^+^-ATPase (VHA). *Brown arrows*, the signalling ABA pathway; ABA (on BL background) activates (→) VHA and inhibits (––I) AHA2, consequently alkalinizing both the cytosol (CYT) and the xylem sap (pH ↑), while acidifying the vacuole (VAC, pH ↓). ABA (independently of BL illumination) activates directly the GORK / SKOR K^+^-efflux channels allowing K^+^ efflux which leaves behind uncompensated negative charges (A^-^), thereby hyperpolarizing the BSC (i.e., lowering their membrane potential, E_M_). H^+^ and K^+^, imported via H^+^/K^+^ and H^+^/NO_3_^-^ symporters (SYM), depolarize the BSCs and acidify their cytosol, in balance with the other transporters. The inclination of arrows directly associated with the transporters indicates the direction of the electrochemical potential gradient for the transported ion. Other details can be found in the text.

Unlike in illuminated BSCs, in the dark, ABA treatment hyperpolarized the BSCs (Fig. 6B), which signified a change in the balance of electrical charges across the plasma membrane in favor of a net influx of negative ions (anions) or a net efflux of positive ions (cations). Could this be due to ABA-induced activation of K^+^ efflux channels (SKOR and / or GORK, the genes of which we found in the BSCs’ transcriptome; Wigoda et al., 2017), and the resulting K^+^ efflux? A similar ABA-induced hyperpolarization, concomitant with H^+^-ATPase inhibition, was observed in Arabidopsis roots and was shown to result from the activation of GORK channels and K^+^ efflux (Planes et al., 2015). Further studies will explore this mechanism in BSCs.

## DISCUSSION

ABA is a central regulator of plant water status and a very large body of research focused on guard cells (GCs) revealed many elements of the ABA signaling cascade in the GCs (e.g., Kollist et al., 2019). Here we focus on ABA effects on the much understudied, ususally difficult-to-access intra-leaf bundle sheath cells (BSCs), and in particular, on the BSCs ABA responses as reflected in their pH (pH_EXT_ and pH_CYT_) and their membrane potential (E_M_).

### The ion fluxes underlying ABA-induced xylem sap alkalinization

Similar to the rise of the XPS pH in the detached leaf due to the inhibition of the BSCs’ plasma membrane AHA by vanadate (Grunwald et al., 2021), the alkalinization of XPS by ABA – the ABA response – signified the inhibition of the BSCs’ AHA by ABA. As before, we attribute this XPS alkalinization to an imbalance of proton fluxes consequent to AHA inhibition, i.e., to a net influx of protons from the xylem to the BSCs (according to the biophysical principles outlined, for example, by Sze and Chanroj, 2018 known as the proton motive force, PMF). This PMF into the BSCs was likely carried by the plasmalemmal H^+^ symporters, such as an H^+^/ K^+^ symporter and an H^+^/NO3^-^ symporter. Notably, the K^+^ uptake permeases, AtKT2 and AtKUP11, are found in the BSCs transcriptome and are likely to be H^+^-coupled (Wigoda et al., 2017; Elumalai et al., 2002; Ahn et al., 2004), as is also the H^+^/NO_3-_ uptake carrier, AtNRT1.1 (Fang et al., 2016; Wigoda et al., 2017). To enhance such potential proton influx we conducted these experiments in an XPS solution containing 1 mM KCl and 10 KNO_3_ mM (as in Grunwald et al., 2021; see Solutions).

### The effect of vanadate on the pH_CYT_ of BSCs protoplasts

We showed previously that petiole-fed vanadate (1 mM) elevated the pH_EXT_ in detached leaves due to inhibition of proton extrusion from the BSCs and the consequent unbalanced leak of protons from the 1 mM KCl+10 mM KNO_3_- containing XPS (Fig. 1 in Grunwald et al., 2021) into the BSCs. However, the pH_CYT_ of the vanadate-treated BSCs protoplasts bathing in 5 mM KCl remained unaffected (Fig. 4B), although both major BSCs’ H^+^-ATPases, AHA1 and AHA2 (Wigoda et al., 2017), are vanadate sensitive, and therefore should have been inhibited, followed by the above-mentioned proton leak imbalance. This unexpected result can be explained, perhaps, by the proton-buffering capacity of the cytosol able to contain the relatively small proton influx into the protoplasts (due, in turn, to the relatively small concentration of the H^+^-cotransport-enabling ions in the 5 KCl bath). In addition, or, alternatively, the small H^+^ influx via the H^+^/K^+^ symport from the low-K^+^ bath could be overcome by vanadate-*insensitive* proton pumps in the cytosol-delimiting membranes. These could be the vacuolar proton pumps, and/ or, the plasma membrane AHA3 which is at least 3-fold less sensitive to vanadate than AHA1 or AHA2 (Palmgren and Christensen, 1994); in support of such compensatory mechanisms we may cite the absence of XPS alkalinization when the xylem perfusate contained vanadate but only 1 mM K^+^ (Grunwald et al., 2021). How are the vanadate-treated BSCs protoplasts different from the BSCs protoplasts of the AHA2-activity-devoid *aha2-4* mutant plant, in which the control (C) pH_CYT_ (6.3) was so much lower than the control WT’s pH_CYT_ (7.7, Fig. 4A)? We remain puzzled at the apparent absence or inactivity of a pH_CYT_-corrective mechanisms in the protoplasts of the mutant.

### Early elements of ABA signaling in BSCs compared to those in guard cells (GCs)

The demonstrations of ABA responses in individual protoplasts – BSCs depolarization (Fig. 3) and their cytosol alkalinization (Figs. 4 and 5) are evidence for the BSCs autonomy regarding ABA signaling in these responses. Could they be initiated in BSCs by ABA via a different route than in GCs? In support of a GCs-resembling BSCs autonomy of early ABA signaling, the transcriptome of BSCs (Wigoda et al., 2017) attests to the expression in BSCs of eight out of 14 Arabidopsis PYLs, six of which participate in ABA-induced stomata closure (Suppl. Table S1; Gonzalez-Guzman et al., 2012, and references therein; Merilo et al., 2015), including PYL2, shown to be sufficient for guard cell responses to ABA (Dittrich et al., 2019).

BSCs express also eight out of the nine Arabidopsis clade A PP2C genes, including the five implicated in early ABA signaling in guard cells (Suppl. Table S1; Wigoda et al., 2017; Merilo et al., 2015). As in stomata, the combinations of these proteins in BSCs is likely to convey ABA-induced signals of varying intensities (Tirsch et al., 2017) and the variety of these proteins in BSCs suggests a rich repertoire of their ABA responses, and a strong possibility of their complete autonomy in ABA signaling.

### The BSCs’ AHA2 is a target of ABA signaling

Notwithstanding the possible similarity between the GCs’ and the BSCs’ initiation of ABA signaling, and based on our earlier demonstration of the AHA2’s prominent role in the leaf xylem sap acidification (Grunwald et al., 2021; Grunwald et al., 2020), we now identify AHA2, rather than AHA1, as the BSCs- specific target of this ABA-inducible PP2C-mediated signaling cascade. This is based on (a) the abolition of the ABA response (Fig. 2B) in the xylem of the detached leaf of the *aha2-4* mutant devoid of AHA2 activity (Haruta et al., 2010; Haruta and Sussman, 2012), and (b) the restoration of the ABA response in the *aha2-4* complemented with the BSCs-directed *SCR:AHA2* Grunwald et al., 2021; Fig. 2C). The complete lack of pH_EXT_ response to ABA in the *aha2-4* mutant (Fig. 2B), suggests, somewhat surprizingly, that in spite of the relatively abundant expression of AHA1 in the BSCs plasma membrane (Wigoda et al., 2017), and unlike the well-documented ABA effect on AHA1 in the guard cells (Merlot et al., 2007) ABA appears not to have affected the BSCs’ AHA1, nor any of the other BSCs plasma membrane AHAs, beside AHA2. Hence, we can conclude that the inhibition by ABA of the BSCs’ AHA2 is both necessary and sufficient for a “normal” (WT-like) level of ABA-induced XPS alkalinization. While this resembles the Arabidopsis seedlings hypocotyl in which ABA inhibited AHA2 (Hayashi et al., 2014) this response differs from the Arabidopsis GCs’, in which ABA inhibited AHA1 (Hayashi et al., 2011).

Moreover, the same mechanism which caused the XPS alkalinization – the net influx into the BSCs of protons combined with an influx of K^+^, both carrying positive charges accross the BSCs plasma membrane – likely underlies the ABA-induced depolarization of the isolated WT BSCs protoplasts (Fig. 3)(Brault et al., 2004). Together, these results attest to the prominent effect of ABA in the BSCs consisting of the inhibition of the BSCs’ AHA2 through the BSCs-autonomous ABA signaling cascade.

Importantly, the ABA responses of the BSCs in the current study reaffirmed the dual role of the BSCs’ AHA2 in: (a) in maintaining the “resting” low pH of the xylem sap (low pH_EXT_) during the day, as demonstrated recently (Grunwald et al., 2021; Grunwald et al., 2020) and (b) in maintaining the BSCs membrane potential in an illuminated leaf at a hyperpolarized range (also as in Grunwald et al., 2020), akin to the dual role of AHA1 in GCs (reviewed in Jezek and Blatt, 2017).

### ABA alkalinizes the BSCs’ cytosol by activating VHA

Cytosolic pH (pH_cyt_) change is a known signal in the ABA signaling pathway; cytosol acidification was demonstrated in Arabidopsis epidermal root cells (using pH-microelectrodes; Planes et al., 2015). In contrast, cytosol alkalinization was demonstrated, among others, in corn coleoptiles and parsley hypocotyls and their roots (using BCECF; Gehring et al., 1990), in guard cells of orchid leaf epidermal strips (using BCECF; Irving et al., 1992), and guard cells of Arabidopsis (using BCECF; (Bak et al., 2013), and in BSCs, in our experiments (using SNARF1, Figs. 4 and 5).

Four results converged to support the VHA activity as responsible for the ABA- induced alkalinization of the BSCs’ cytosol: (a) the cytosol alkalinization by ABA of a*ha2-4* mutant devoid of AHA2 activity (Fig. 4A) eliminated the AHA2 as a culprit; (b) the general P-type ATPases inhibitor, vanadate, eliminated most of the plasma membrane proton pumps (including AHA2 and AHA1) as contributing to ABA-induced alkalinization (Fig. 4B), leaving the vacuolar H^+^-pumps and perhaps also AHA3; (c) the abolition of ABA-induced alkalinization by the VHA-specific (Muroi et al., 1994) inhibitor, bafilomycin A1 (Fig. 5A) and (d) the abolition of ABA-induced cytosol alkalinization in the double mutant *vha-a2 vha- a3,* devoid of VHA activity (Krebs et al., 2010, Fig. 5B).

Arabidopsis possesses at least three vacuolar proton pumps: the multi-subunit V- type H^+^-ATPase, VHA (Schumacher and Krebs, 2010), the homo-dimeric V-PPase, AVP1 (AT1G15690; Segami et al., 2018), and the P-type group III H^+^- ATPase, AHA10 (AT1G17260; Appelhagen et al., 2015). Among these, our results singled out the VHA as solely responsible for the rapid (1-15 min) ABA-induced cytosol alkalinization.

That the vacuolar H^+^-ATPase could be stimulated by ABA and was then capable of alkalinizing the cytosol could be deduced from early experiments which focused on the increased activity of the vacuolar H^+^-ATPase induced by prolonged ABA treatments (days) and demonstrable as the acidification of vacuolar vesicles from mature leaves of *Mesembrianthemum crystallinum* (Barkla et al., 1995), or of microsomal vesicles from Arabidopsis etiolated seedlings (Krebs et al., 2010). Relatively brief ABA treatments (30 min), more similar to ours, induced vacuole acidification and concomitant cytosol alkalinization in guard cells of *Vicia faba* and of Arabidopsis (Bak et al., 2013), resulting, however, from ABA-induced activation of *both* the VHA and the AVP1 (VHP1). This was concluded based on mutants of both pumps (respectively, *vha-a2 vha- a3* and *vhp1*); and the requirement for both intact proton pumps together for a normal ABA-induced stomata closure in mature Arabidopsis rosettes (Bak et al., 2013). In contrast, the constitutive overexpression of AVP1 in the severely dwarfed *vha-a2 vha-a3* mutant background *did not* increase the mature rosette size, *neither did it* correct the overly elevated leaf cell sap pH (representing mainly the vacuolar sap pH), or the seedling root vacuolar pH (Kriegel et al., 2015). This ineffectiveness demonstrated the overall inability of the vacuolar PPase to compensate for the absence of the vacuolar ATPase activity, in spite of its (the PPase’s) theoretical and demonstrable capability to contribute to the vacuole acidification in the guard cells (Kriegel et al., 2015; Segami et al., 2018). Our own results, showing that a selective elimination of the VHA pump activity – either by bafilomycin A1 or by the double mutation – abolished completely the ABA-induced elevation of the BSCs’ pH_CYT_ (Fig. 5), suggest that, in contrast to GCs, the BSCs’ VHA alone – and not their AVP1 – appears to be activated as a target of the ABA signaling pathway inducing cytosol alkalinization.

### A role for additional tonoplast transporters in BSCs’, pH_CYT_ maintenance?

What underlies the difference between the highly elevated pH_CYT_, about 7.9, in the *un*treated (control, C) *vha-a2 vha-a3* double mutant (Fig. 5B) – as compared to about 7.2 in the *un*treated (C) WT BSCs (Figs. 5A, 5B)? A few explanations can be suggested: (i) the pH_CYT_ of the untreated (C) *vha* mutant was elevated via the activity of plasma membrane AHAs other than AHA2, since AHA2 is reportedly almost inactive at pH_CYT_ above neutral (Hoffmann et al., 2019); (ii) a compensatory H^+^ pumping by AVP1 (the vacuolar PPase), suggestedly responsible for the mutant viability (Krebs et al., 2010; Sze and Chanroj, 2018), could have over-alkalinized the cytosol. However, the *vha* mutant’s vacuolar pH (pH_VAC_) remained about 0.5 units above the WT’s pH_VAC_ of 5.8 (Krebs et al., 2010), not what would be expected from a vigorous H^+^ pumping into the vacuole by the AVP1. Thus, (iii) a possible explanation for both pH_CYT_ and pH_VAC_ would be a combination of the two aforementioned pumping mechanisms (across both cytosol boundaries), combined with enlarged H^+^ efflux from the vacuole into the cytosol via proton exchangers of the tonoplast – such as the H^+^/K^+^ exchangers NHX1 and NHX2 (Andrés et al., 2014) or the H^+^/anion CLC exchanger (De Angeli et al., 2006).

Can the same be argued to explain also the difference between the pH_CYT_ of the ABA-treated *vha* mutant (approx. 8.1, Fig. 5B) and the pH_CYT_ of the ABA + bafilomycin A1-treated WT (7.2, Fig. 5A), i.e., in a situation with *both* AHA2 and VHA inactive?

We propose that the lack of effect of ABA treatment on the *vha* mutant’s high pH_CYT_ (approx. 8.1; Fig. 5B), is due to the mutant BSCs’ lack of any ABA- susceptible activity of H^+^-pumps VHA and AHA2 (the latter, due to the abovementioned inactivity at pH_CYT_ above neutral; Hoffmann et al., 2019). We then propose that the cytosol alkalinization seen under the brief treatment of the WT BSCs with bafilomycin A1 alone (pH_CYT_ 7.2 to 7.6, Fig. 5A), was due to the inhibition of the background VHA activity, which, in turn altered the pH_VAC_, activating / or enhancing, in compensation, the vacuolar AHA10 activity which pumped protons from the cytosol into the vacuole. In contrast, the relatively low pH_CYT_ seen when ABA was added together with bafilomycin A1 (the same as in control, 7.2; Fig. 5A), the alkalinizing effect of AHA10 activity resulting from VHA inhibition may have been abolished byABA-induced inhibition of AHA10, similar to the ABA-induced inhibition of AHA2. Cytosol acidification, on the other hand, may have been prevented by the buffering capacity of the cytosol containing the slow leak of protons from the bath and from the vacuole via the cotransporters/exchangers, at least within the short time window of our experiment. Clearly, detailed studies are needed to test these hypotheses.

### Physiological significance

Since pH_EXT_ and pH_CYT_ have been shown to affect the activities of aquaporins and ion transporters, the ABA-induced alkalization within and outside the BSCs offers a mechanism underlying a possible effect of drought (mediated by ABA) on the BSCs function as a selective barrier to ion and water radial transport into the leaves. Such a mechanism – modifying the BScs’ selective barrier function by altering the pH in the membrane vicinity – could be in operation also under other stresses signaled by ABA – such as cold or salinity. Consequently, our results highlight both proton pumps (AHA2 and VHA) in the BSCs as attractive targets for future designs of stresses-tolerant plants.

## MATERIALS AND METHODS

### Plant Material

#### Genotypes

We used two accessions of wild type (WT) *Arabidopsis thaliana*: Landsberg *erecta* (Ler) and Columbia (Col), and a few mutants and transformants: the whole plant PP2C (*ABI1,* At4G26080.1) knockout, *abi1-1* (in Ler background; Koornneef et al., 1984), obtained from TAIR, stock CS22), *abi1- 1* (in Col background; aka *abi1-1C*; Umezawa et al., 2009), a kind gift from the lab of Dr. K. Shinozaki); *SCR*:*abi1-1* (Col) plants, i.e., WT(Col) plants transformed with the *abi1-1* gene under the BSCs-directing Scarecrow (SCR) promoter (Wysocka-Diller et al., 2000; and see below); *aha2-4* (Col) (SALK_ 082786), a whole-plant T-DNA insertion-knockdown of the H^+^-ATPase, *AHA2* (Haruta et al., 2010; Haruta and Sussman, 2012), obtained from the Arabidopsis Biological Resource Center (Ohio State University); *aha2-4* plants complemented with the BSCs-directed *SCR:AHA2* (Grunwald et al., 2021), and *vha-a2 vha-a3* (Col), the double mutant of the two tonoplast-localized isoforms of the A subunit of VHA (Krebs et al., 2010); a kind gift from Dr. Schumaker’s lab).

In addition, single-cell experiments employed GFP-labeled BSCs protoplasts from the following plant genotypes harboring SCR:GFP: WT (Ler; Shatil-Cohen et al., 2011), WT (Col), *aha2-4* (Col) and *vha-a2 vha-a3* (Col). We used plants from progressively higher generation numbers selected for brighter GFP labeling (Attia et al., 2020) of BSCs.

#### Plant growth

The Arabidopsis plants were grown in a growth chamber as detailed previously by Grunwald et al., 2021.

### SCR:abi1-1 (Col) constructs and plant transformation

For construct assembly, the MultiSite Gateway Three-Fragment Vector Construction Kit (Invitrogen) was used according to the manufacturer’s instructions. Bundle-sheath-ABA-insensitive (*SCR*:*abi1-1*) plants were constructed similarly to *fa* plants of (Negin et al., 2019). Briefly, the *abi1-1* gene (Koornneef et al., 1984) and the SCR promoter were cloned into pDONR plasmids and then inserted into the binary pB7M24GW (Invitrogen) plasmid. The plasmids were inserted into Arabidopsis WT (Col) plants using the floral-dip method (Clough and Bent, 1998). The presence of the transgene containing the G to A base substitution was confirmed by sequencing. The validation study (Suppl. Fig. S2) was performed on three independent lines of each construct, using homozygous t3 and t4 lines.

Quantification of *abi1-1* expression in WT, *abi1-1* and *SCR:abi1-1* plants by restriction enzymes was as described by (Negin et al., 2019). Briefly, total RNA was extracted from leaves and cDNA was prepared. The ABI1_174F and ABI1_1064R primers (ABI1_174F: TCTGGGTCACATGGTTCTGA, ABI1_1064R: CCATCTCACACGCTTCTTCA) were used for amplification of both ABI1 and *abi1-1*. The amplified 911bp fragment was cleaned and 1000 ng of the amplified fragment was digested for 2 hours by NcoI and loaded onto 1% agarose gel. The gel was photographed under ultraviolet (UV) light and digital image analysis was performed using ImageJ.

***Stomatal aperture*** was measured as described by (Yaaran et al., 2019). Briefly, epidermal peels were soaked in a ‘closure-enabling’ solution (ibid.) under a light intensity of ∼150 μmol m^−2^ s^−1^. After 1.5 h, ABA ((+)-cis, trans abscisic acid; Biosynth; Staad, Switzerland) was added to a final concentration of 10 μM. DMSO at the same concentration was added to the control. All stomata were photographed under a bright-field inverted microscope (1M7100; Zeiss; Jena, Germany) on which a Hitachi HV-D30 CCD camera (Hitachi; Tokyo, Japan) was mounted. Stomatal images were analyzed to determine aperture size using the ImageJ software (http://rsb.info.nih.gov/ij/).

### Fluorescence imaging

#### Fluorescent dyes for the determination of H^+^-Pumps activities

We monitored the pH established as a result of manipulations in the H^+^-pumps activities using fluorescence imaging of pH- and membrane potential (E_M_)- reporting ratiometric fluorescent probes: (a) pH_EXT_, the pH of the xylem perfusion solution (XPS), using FITC-dextran, FITC-D (fluorescein isothiocyanate conjugated to 10 kD dextran, a dual-excitation pH probe, (Grunwald et al., 2021) (b) pH_CYT_, the BSCs’ cytosolic pH – using SNARF1 (5-(and 6)-carboxy SNARF-1 acetomethyl ester acetate), a dual emission pH probe, preloaded into the isolated protoplasts (Sano et al., 2010), and (c) the BSCs’ E_M_ – with di-8-ANEPPS (4-(2- [6-(Dioctylamino)-2-naphthalenyl]ethenyl)-1-(3-sulfopropyl)pyridinium inner salt), a dual excitation potentiometric probe (Pucihar et al., 2009; Wigoda et al., 2017; Grunwald et al., 2020), also preloaded into the isolated protoplasts.

#### pH_EXT_: detached leaves preparation and imaging

The experiments were performed between 9 AM and 1PM. In the morning of the experiment, shortly before the lights went on in the growth chamber (at 9 AM), several 6-7 week old plants plants were placed in a nearby dark room at roughly the same temperature. Leaves, approximately 2.5 cm long and 1 cm wide, were excised under green light with a sharp blade at 20 min intervals, placed in Eppendorf vials with unbuffered xylem perfusion solution (XPS, see Solutions) containing 100 μM of FITC-D, added from a 10 mM stock in DDW, as described by Grunwald et al., 2021, without or with the addition of 10 μM ABA. The vials (wrapped in aluminium foil) were placed in humidity boxes (Grunwald et al., 2020), and returned to the growth room for 1.5 -2 hours in full light. Each leaf was then brought back individually for a microscope slide preparation, which lasted about 2 min, it remained on the microscope stage for about 5 min. and during these 7 min it remained under dark (<0.025 µmol m^-2^ s^-1^ between 400 and 850 nm, as determined using a spectrometer (Maya2000-Pro with the 400 m head, from Ocean Optics, Germany, www.oceanoptics.eu).

The veins of detached leaves were imaged using *setup I* (based on an inverted microscope Olympus-IX8, as detailed by (Grunwald et al., 2021). Image capture and image analysis of the intra-xylem pH (pH_EXT_), as well as pH_EXT_ calibration were exactly as already described (ibid.), except a new conversion relationship (Suppl. Fig. 1) was used in the current analyses.

#### pH_CYT_: protoplasts preparation and imaging

The experiments were performed between 10 AM and 4 PM. BSC protoplasts were isolated from 6-8 weeks old *SCR:GFP*-labeled plants in an approximately 30 min-long procedure, as described (Shatil-Cohen et al., 2014) and kept in an Eppendorf vial at room temperature (22 ± 2 °C) under DL, as above, until use. The protoplasts were imaged in *setup II*, consisting of an inverted epifluorescence microscope (Eclipse Ti-S, Nikon, Tokyo, Japan) coupled to an IXON Ultra 888 camera (Andor, UK) via an OptoSplit device (Cairn research, UK). A Nikon 40X objective (Plan Fluor 40x/ 0.75, OFN25 DIC M/N2) was used for protoplast viewing and imaging. The fluorescence excitation beam was delivered by a xenon lamp monochromator (Polychrome II, Till Photonics, Munich, Germany), under the control of IW6.1 (Imaging Workbench 6.1) software (Indec Biosystems, Santa Clara, CA). All excitation and emission filters were from Chroma Technology Corp. (Bellows Falls, VT, USA).

400 µL of bath solution which included 50 µL of protoplasts suspension, SNARF1-AM (5µM final conc., see Solutions) and pluronic acid (1 % final conc., see Solutions), premixed in an Eppendorf vial, were added to the experimental chamber and incubated for 30 min at RT. While the cells settled and stuck to the chamber glass bottom, the membrane-permeant SNARF-AM was digested by cytosolic hydrolases releasing SNARF1, the membrane impermeant free fluorescent ionic form of the dye which largely remained trapped in the cytosol – the most alkaline cellular compartment – for the duration of the recording.

At min 20 into SNARF1-AM incubation, blue and red (BL+RL) illumination was aplied for 10 min (from a home-made illuminator of crystal clear LEDs from Tal- Mir electronics, Israel), consisting of RL, 660 nm, roughly 195 µmol m^-2^ s^-1^ and BL, 450 nm, roughly 25 µmol m^-2^ s^-1^ at the protoplast level. During the last 1 min of illumination, the cells were flushed to remove the external SNARF-AM, with bath solution without or with 3 µM ABA (the flush volume, ≥ 5 mL, was more than 10X the chamber volume of ∼400 µL). Thereafter, the RL+BL illuminator was switched off. An individual, perfectly round BSC (diameter of 25-32.5 µm) was selected for pH_CYT_ imaging based on its GFP fluorescence (Ex.: 490/15 nm, Em.: dichroic mirror: 515 nm and band-pass barrier filter: 525/50 nm). The selected protoplast was focused at its largest diameter under green-filtered phase contrast illumination (‘transmitted light’) and its image was recorded. Immediately thereafter, the SNARF1 within the protoplasts was excited by 550/15 nm 50 ms long light pulse, the emitted fluorescence was deflected by a 570 nm dichroic mirror (T570 LPXR) into the OptoSplit device, further filtered via a 575 nm long-pass filter (ET575LP), then split by a 612 nm dichroic mirror (T612 LPXR-UF2) into two beams, filtered in parallel: via a 640/20 nm band-pass filter (ET640/20M) and via a 585/20 nm band-pass filter (ET585/20M), and the two images were recorded simultaneously on two separate halves of the camera CCD sensor. Then, a next BSC cell was selected in the same chamber, and so on, until 6 min elapsed since the bath flush with the SNARF1-AM-free solution (±ABA). A next batch of cells was then transferred to a clean chamber for further imaging.

### Image analysis

The emitted fluorescence intensities in the recorded images were evaluated using FIJI (Abràmoff et al., 2004; Schindelin et al., 2012). The measure of the cytosolic pH was the ratio between the fluorescence intensity at 640 nm which *in*creases with pH, and the fluorescence intensity at 585 nm which *de*creases with increasing pH (Sano et al., 2010), each corrected for the corresponding average background fluorescence. To minimize possible errors due to a possibly imperfect overlap of the two recorded images in each pair, the ratiometric analysis was performed on the *averaged* intensities of all the pixels in the image which passed the selection criteria. The selection criteria were applied to the usually brighter image at 640 nm; we excluded manually those image regions which were selected automatically but departed obviously from the circumference ring and regions which appeared saturated (the selected areas are enclosed within yellow lines in the Suppl. Fig. S3A).

#### pH_CYT_ calibration using SNARF1

To calibrate the SNARF1 emission ratio values vs cytosolic pH (pH_CYT_) values, we used SNARF1-loaded WT (Ler) protoplasts in solutions containing either 200 mM K^+^ (as in Fig. S4B) or 250 mM K^+^ (as in Fig. S4C) (see Solutions below). The incubation with SNARF and the imaging were conducted as described above except (a) there was no BL+RL illumination, (b) the flushed-in bath solution contained 5 µM nigericin (an H^+^/K^+^ exchanger) and was bufferred to different pHs (pH_EXT_; see Solutions) and (c) the protoplasts were imaged within the first 3-5 min from the nigericin flush-in, before it reached the endomembranes and equilibrated the cytosol with other compartments.

Assuming that an equilibrium is attained within a few minutes (Thomas et al., 1979), proton concentrations ratio across the plasma membrane becomes equal to the K^+^ concentrations ratio:

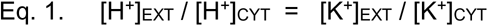

and the internal pH (pH_CYT_) becomes equal to the external pH (pH_EXT_) when [K^+^]_EXT_ = [K^+^]_CYT_.

*Potential errors sources of SNARF1 calibration:* Because the [K^+^]_CYT_ in BSCs in our experiments is not known, the resulting deviation of pH_CYT_ from the imposed pH_EXT_ value is also unknown. Thus, if [K^+^]_EXT_ that we use > the true [K^+^]_CYT_, then the *true* pH_CYT_ that we impose > pH_EXT_ which we *think* that we impose. Thus, for example (based on a simulation using Eq. 1), if we use a calibration solution with 200 mM K^+^ at pH 6.5 (i.e., [K^+^]_EXT_ = 200 mM, pH_EXT_ = 6.5), while the true (but unknown) [K^+^]_CYT_ = 400 mM, the resulting (also unknown) pH_CYT_ will be 6.2, rather than 6.5, but if the the true [K^+^]_CYT_ = 100 mM, the resulting pH_CYT_ will be 6.8, rather than 6.5, also unbeknown to us. These potential errors should affect mainly the *absolute* pH values, but the relative pH changes still remain valid.

Moreover, inexplicably, WT (Col) Arabidopsis were recalcitrant to the calibration with 5 µM nigericin, which was therefore performed only on WT (Ler) protoplasts and we assumed its validity also for the other genotypes.

### E_M_: protoplasts imaging

The protoplasts were imaged in the Nikon-microscope-based *setup II*, using Di- 8-ANEPPS, as described earlier by Grunwald et al., 2021, with a few changes. A 50 µL drop with leaf protoplasts was placed atop a glass bottom of an experimental chamber, allowed to settle and stick to the glass for 10 min, then 400 µL isotonic bath solution containing 30 µM ANEPPS and 1% pluronic acid, without or with 3 µM ABA, was gently added to the chamber, filling it completely. Then, blue and red (BL+RL) light was turned on for 10 min then turned off. A BSC protoplast, selected based on its GFP fluorescence, was focused at its largest diameter and its transmitted-light image was recorded as above; then it was exposed to a pair of consecutive 3 ms-apart, 50 ms-long excitation pulses, of 438 nm, then 531 nm. The di-8-ANEPPS fluorescence emitted from the membrane (the dye which remained dissolved in the bath did not fluoresce) was filtered via a dichroic mirror of 570 nm and 585/20 nm emission band-pass filter and the pair of the resulting consecutive images was recorded by the camera. A next BSC cell was then selected in the same chamber, and so on, for about 5 min, until 15 min elapsed since di-8-ANEPPS (+/ABA) addition. A next batch of cells was then transferred to a clean chamber for further imaging.

***Image analysis and fluorescence ratio calibration*** of di-8-ANEPPS were as detailed earlier (Grunwald et al., 2020). Briefly, we obtained the following positive relationship between the membrane potential, E_M_, imposed using patch-clamp, and the mean fluorescence ratio (F-ratio) of the cell:

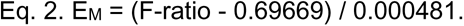

Importantly, while this quantitative conversion could not be applied to the data of Fig. 6 due to an unexpected replacement of the excitation source, we deem it sufficient to rely on the established positive correlation of F-ratio vs. E_M_ to use the change in the F-ratio between treatments as a qualitative indication of the relative *direction* of the treatment-induced E_M_ change: depolarization or hyperpolarization.

### Solutions

***XPS, xylem perfusion solution*** (in mM): 10 KNO_3_, 1 KCl, 0.3 CaCl_2_ and 20 D-sorbitol. Upon preparation, when unbuffered, the pH of this solution was 5.6 - 5.8 and its osmolarity was approx. 23 mOsm.

Bath solution for SNARF1 (pH_CYT_) calibration (in mM):

- *<u>200 mM K^+^</u>*, as in Suppl. Fig. S4B (in mM): 200 K-Gluconate, 1 CaCl_2_, 4 MgCl_2_, 10 HEPES, different pHs (pH 6.0, 6.5,7,7.5,8 and 8.5) were adjusted using 1 M NMG, osmolality: 435 mOsm, adjusted w/ D-sorbitol;
- *<u>250 mM K^+^</u>*, as in Suppl. Fig.S4C (in mM): 250 K-Gluconate, 1 CaCl_2_, 4 MgCl_2_, 10 HEPES, different pHs (pH 6.5,7,7.5,8 and 8.5) were adjusted using 1 M NMG, osmolality: 456 mOsm.

***Bath solution for protoplasts’ pH_CYT_ determination*** (in mM): 5 KCl, 1 CaCl_2_, 4 MgCl_2_, 10 MES; pH 5.6; osmolality: 435 mOsm, adjusted w/ D-sorbitol.

**Pipette solution for di-8-ANEPPS (E_M_) calibration** (in mM): 112 K-Gluconate, 28 KCl, 2 MgCl_2_, 10 HEPES, 328 mOsm Sorbitol, pH 7.5, Osmolarity: 480 mOsm

***Bath solution for protoplasts’ E_M_ determination*** (in mM): 5 KCl, 1 CaCl_2_, 4 MgCl_2_, 10 MES; pH 5.6; osmolality: 435 mOsm, adjusted w/ D-sorbitol.

***ABA:*** (±)-Abscisic acid (Sigma-Aldrich, Cat. # D1049); a 100 mM stock solution in 1 M KOH in DDW was stored in aliquots, protected from light at -20°C.

***FITC-D*:** Fluorescein isothiocyanate–dextran (average mol wt 10,000; Sigma Israel, cat. #: FD10S); a 10 mM stock solution in DDW was stored in aliquots, protected from light, at -20°C.

***Di-8-ANEPPS:*** (Molecular Probes, Life Technologies, cat. # D3167, Oregon, USA), a 10 mM stock solution in DMSO was stored in 10 µL aliquots at -20 °C. ***SNARF1-AM:*** (Molecular Probes/Thermo Fisher cat. #: C-1272); 50 µg portions were dissolved in DMSO sequentially, as needed; a 10 mM stock solution was stored in aliquots at -20°C.

*Pluronic F-127:* (Molecular Probes, Life Technologies, cat. # P6867, Oregon, USA), a 20% stock solution in DMSO was stored in 100 µL aliquots at RT.

***Vanadate:*** sodium orthovanadate, Na_3_VO_4_ (BHD Chemicals Ltd., cat. #30194); the stock solution of 200 mM in DDW was depolymerized by boiling as described in the Sigma-Aldrich Product Information sheet for sodium orthovanadate: https://www.sigmaaldrich.com/content/dam/sigmaaldrich/docs/Sigma/Product_Information_aldrich/docs/Sigma/Product_Information_Sheet/1/s6508pis.pdf.

The stock solution was aliquoted and stored at -20 °C.

***Bafilomycin A1*:** a VHA inhibitor (Sigma-Aldrich, Cat# B-1793); a 0.16 mM stock in DMSO and stored in aliquots, protected from light at -20 °C.

***Nigericin free acid:*** a K^+^/H^+^ ionophore (Molecular Probes/Thermo Fisher cat. #: N-1495); a 5 mg/mL stock in EtOH (J.T. Baker Cat# 64-17-5, type Absolute, Analytical grade), stored in aliquots at -20°C.

## ACKNOWLEDGEMENT

This research was supported by ISF (the Israel Science Foundation, Grant No. 1312/12 to NM and grant No. 1043/20 to MM) and the Ministry of Agriculture, Israel (the Office of the Chief Scientist, grant No. 12-01-0007 to NM). The authors are grateful to Drs. M. Krebs & K. Schumaker for their gift of the double VHA mutant, *vha-a2 vha-a3*, to Dr. K. Shinozaki for the gift of the mutant *abi1-1* (Col), to Ms. V. Schebtaev for the technical lab assistance and to Dr. Dizza Bursztyn of the Hebrew University of Jerusalem for advice on the statistics.

## SUPPLEMENTAL MATERIALS

### Supplemental Table S1

Genes initiating the Arabidopsis ABA signaling pathway found in the BSCs transcriptome.

### Supplemental figures

***FIGURE S1.*** Ratiometric pH calibration of the xylem perfusion solution (XPS).

***FIGURE S2.*** A protein phosphatase 2C (PP2C, ABI1) mediates the ABA-induced alkalinization of the xylem perfusate in the minor veins of Arabidopsis detached leaves.

***FIGURE S3.*** Validation of *SCR::abi1-1* transformation.

***FIGURE S4.*** Ratiometric calibration of cytosolic pH (pH_CYT_) using the dual- emission dye, SNARF1.

