## Supplementary data for "Stress is basic: ABA alkalinizes both the xylem sap and the cytosol of Arabidopsis vascular bundle sheath cells by inhibiting their P-type H^+^-ATPase and stimulating their V-type H^+^-ATPase"

### SUPPLEMENTAL MATERIALS

**Supplemental Table S1**

***Genes initiating the Arabidopsis ABA signaling pathway found in the BSCs transcriptome*** (Wigoda et al., 2017). Arabidopsis BSCs express eight PYR/PYL receptors out of the 14 family members found in Arabidopsis and eight PP2C Clade A phosphatases out of the 9 subfamily members found in Arabidopsis; asterisks mark those expressed in Arabidopsis guard cells (Fig. 2 of Merilo et al., 2015). Note, a review by Cotelle and Leonhardt (2019) lists only four PP2Cs in Arabidopsis guard cells (ABI1, ABI2, PP2CA and HAB1; Cotelle and Leonhardt, 2019).

<u>PYR/PYL ABA receptors</u>

PYR1*, AT4G17870

PYL1*, AT5G46790

PYL2*, AT2G26040

PYL3, AT1G73000

PYL4*, AT2G38310

PYL5*, AT5G05440

PYL6*, AT2G40330

PYL7, AT4G01026

<u>PP2Cs, clade A protein phosphatases 2C</u>

ABI1*, AT4G26080

ABI2*, AT5G57050

HAB1*, AT1G72770

HAB2, AT1G17550

AHG1, AT5G51760

PP2CA*, AT3G11410

HAI1*, AT5G59220

HAI2, AT1G07430

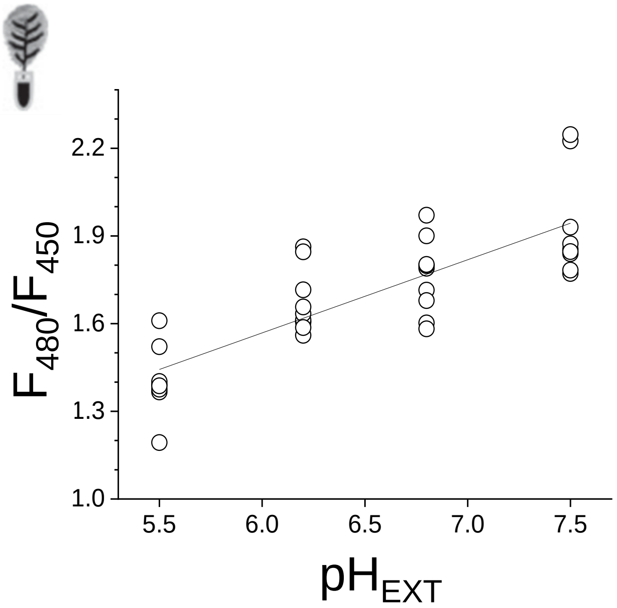


**Suppl. FIGURE S1. Ratiometric calibration of the xylem perfusion solution (XPS) pH.** *In-situ* calibration in detached leaf veins using the membrane-impermeant, dual-excitation pH probe FITC-D (Em: 520 nm; $Ex_1$: 450 nm, $Ex_2$: 480 nm; Materials and methods). WT (Col) leaves were petiole-fed with XPS solution buffered to a series of pH ($pH_{EXT}$) values, as indicated (see Solutions), containing 10 μM FITC-D. Each symbol (a biological repeat) is a mean of 5 individual determinations (technical repeats) per a single leaf. The fluorescence ratio (F-ratio, $F_{480}/F_{450}$) values were fitted linearly with: F-ratio = 0.25029 * $pH_{EXT}$ + 0.06658 (Pearson's R = 0.80516), yielding the conversion relationship: $pH_{EXT}$ = (F-ratio - 0.06658) / 0.25029. Inset: a detached leaf schematic.

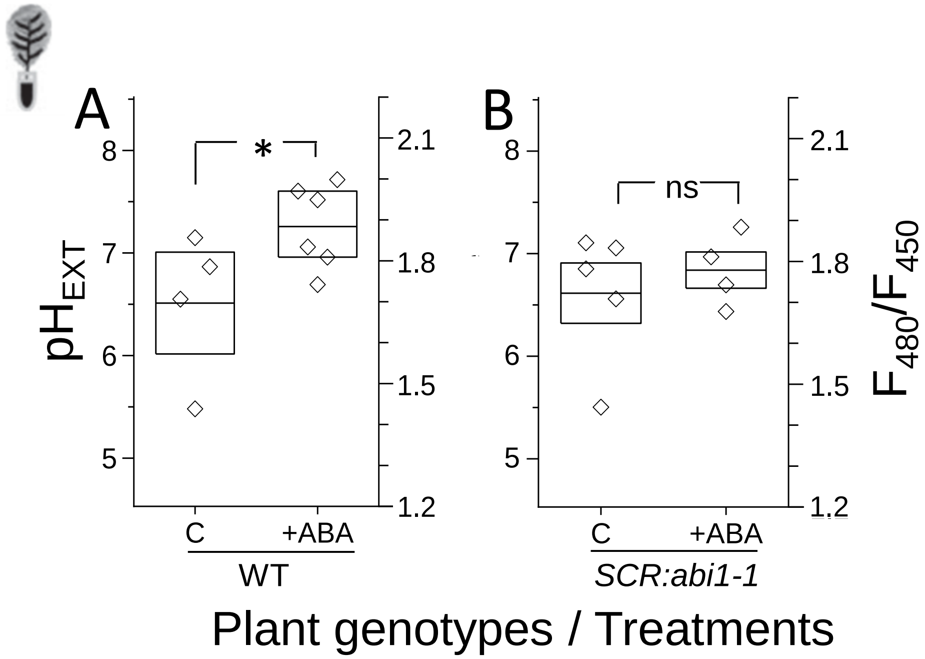


**Suppl. FIGURE S2. A protein phosphatase 2C (PP2C), ABI1, mediates the ABA-induced alkalinization of the xylem perfusate (XPS) in the minor veins of Arabidopsis detached leaves**. **A:** The effect of ABA on the XPS pH ($pH_{EXT}$) in WT *Arabidopsis thaliana*, accession Columbia (Col). Individual pH values (symbols) of non-buffered XPS, each representing a leaf (biological repeat), derived from pixel-by-pixel fluorescence-ratio (F-ratio, $F_{480}/F_{450}$) images of FITC-D (as in Fig. 1 of Grunwald et al., 2020; Materials and methods) and their means ±SE (midline & box; Materials and methods). The data are from two independent experiments. The F-ratio values were converted to pH values based on the *in-situ* calibration curve and the conversion relationship of Suppl. Fig. 1. Asterisk: significant difference (at P= 0.03484, by 1-tailed t-test, equal variance). **B:** Absence of effect of ABA on $pH_{EXT}$ of WT (Col) Arabidopsis transformed with the mutated *abi1-1* (*ABA insensitive*) gene directed to the bundle sheath cells by Scarecrow (*SCR*) promoter. See the validation of the *SCR:abi1-1* transformation in Suppl. Fig. S2. Other details as in A. Note the absence of ABA effect on $pH_{EXT}$ in the *SCR:abi1-1* plants devoid of the phosphatase ABI1 activity in their BSCs. Inset: a detached leaf schematic.

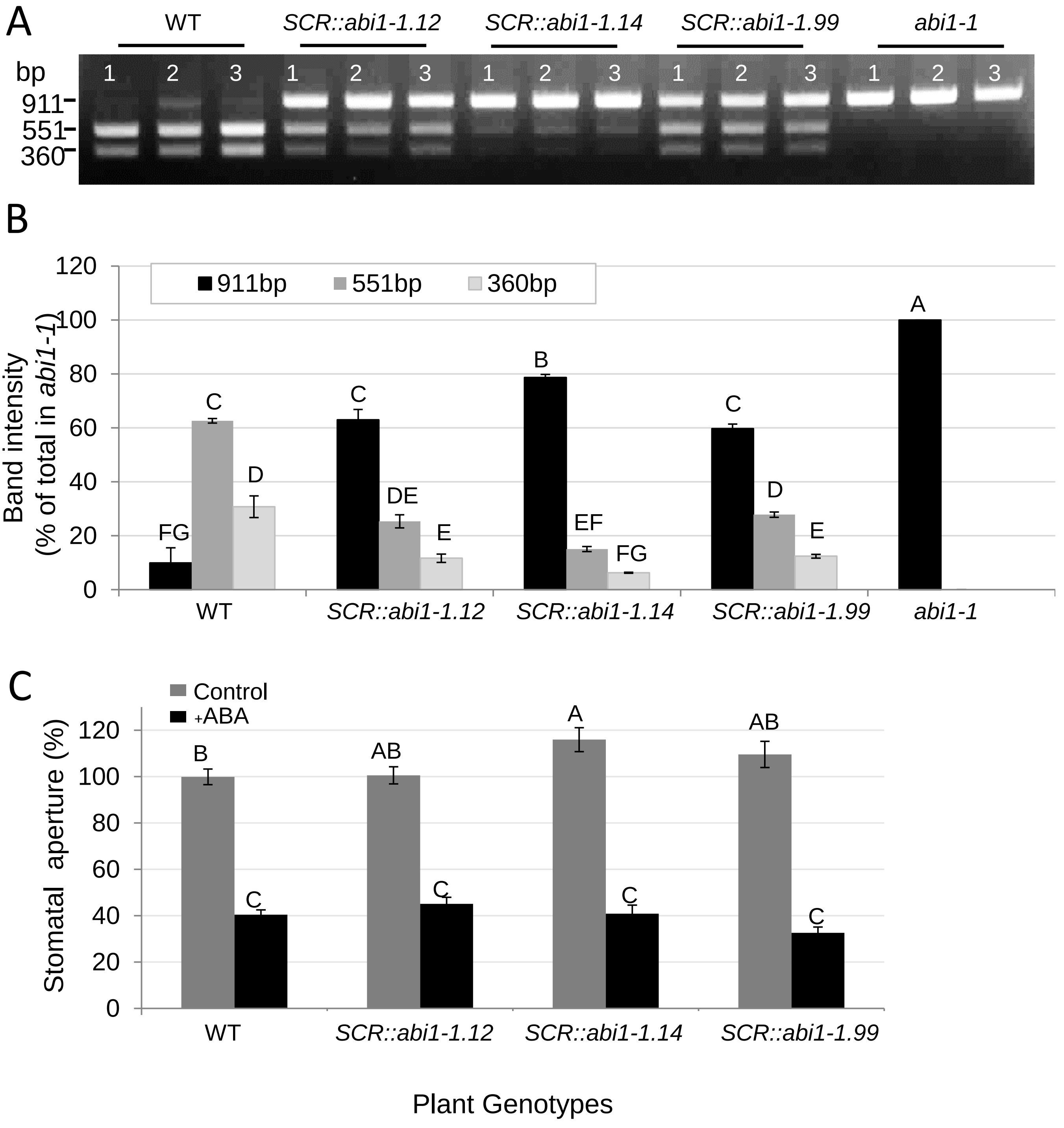


Cont. on next slide

Suppl. Fig. S3, ABA on BSCs pumps, TT. et al., 2021

Cont. from previous slide

**FIGURE S3. Validation of *SCR::abi1-1* transformation. A-B.** Verification by semi-quantitative PCR following digestion by NocI restriction enzyme (Materials and methods). **A.** A representative electrophoresis gel of PCR-amplified *abi1-1* gene (with the mutated NocI restriction site, CCATGG/A, Negin et al., 2019) and thus comprised of an undigested, full-length fragment of 911 base pairs (bp)) and the native (WT) ABI1 gene (yielding the resulting 551bp and 360bp fragments, when fully digested). The plant material consisted of leaves of WT (Col) Arabidopsis, the transformant lines *SCR::abi1-1* (Materials and methods) and the whole-plant mutant *abi1-1* (Col). Note the varying relative amounts of the undigested vs. digested fragments, attesting to different proportions of the negative-dominant *abi1-1* mutant gene expression in the transformed lines. **B.** The intensities of gel bands – as in A – were determined using ImageJ (Materials and methods) for each plant separately and presented as the mean percentage of the given band intensity out of the sum of intensities of all the bands in the lane. Note that the highest proportion of the mutated abi1-1 gene (911 bp) is found in the *SCR::abi1-1*.14 line. **C.** Stomatal response assayed in epidermal peels exposed directly to the ‘closure-enabling’ solution (Materials and methods) without (control) or with 10 μM ABA (+ABA). Results are mean values (± SE, from >90 stomata, from >6 leaves, from 6 plants of each line, from 3 independent experiments, relative to the mean aperture (1.82±0.06 μm) of the WT (Col). Different letters indicate a significant difference according to the Tukey-Kramer test ($P < 0.05$). Note, that plants expressing the ABA-signaling-disrupting *abi1-1* mutation directed to the BSCs under the SCR promoter, retain stomata sensitivity to external ABA.

Cont. form prev. page

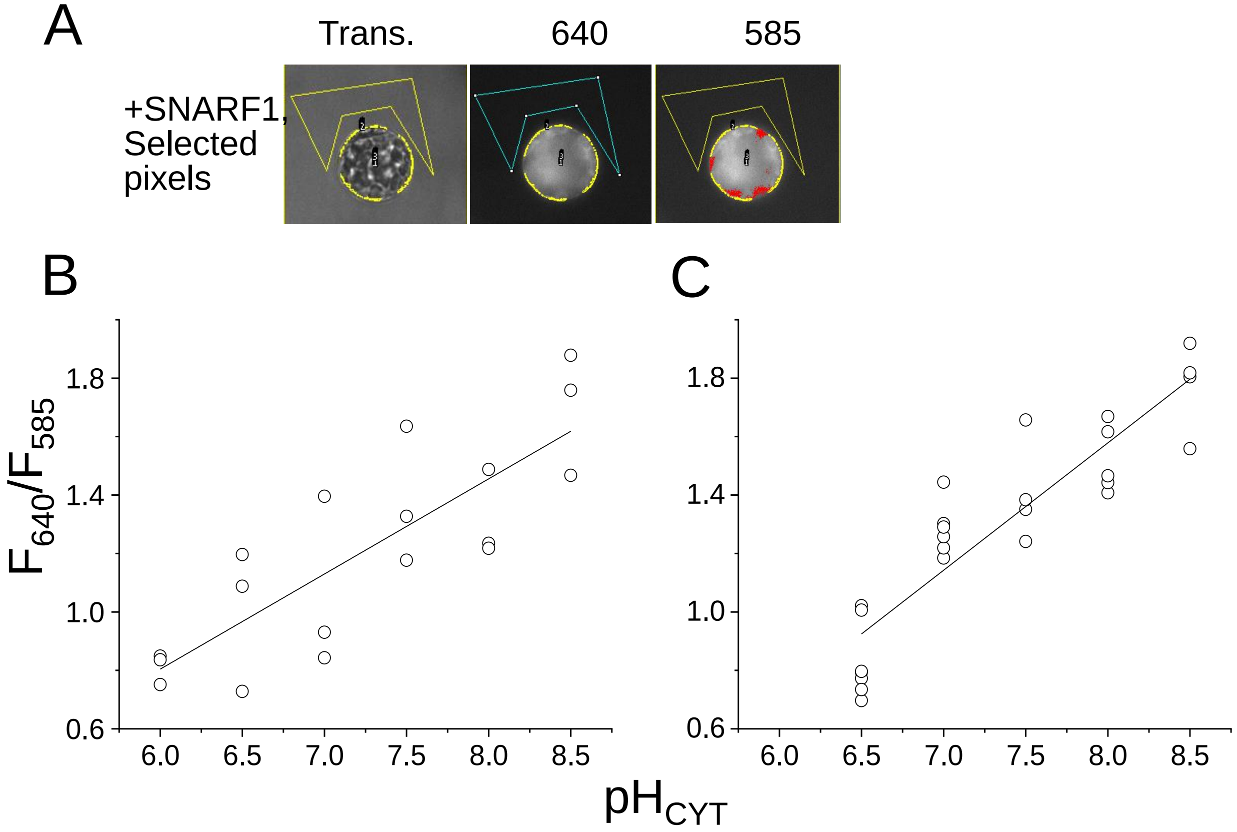


**FIGURE S4. Ratiometric calibration of cytosolic pH ($pH_{CYT}$) using the dual-emission dye, SNARF1.**

**A.** Transmitted light (Trans) and fluorescence images of a BSC protoplast (isolated based on its GFP fluorescence) at the indicated wavelengths, without (-) or after incubation with (+) the ratiometric, dual-emission, pH probe SNARF1 (Ex: 550 nm, $Em_1$:640 nm, $Em_2$: 585 nm; Materials and methods; the autofluorescence of a BSC protoplast devoid of SNARF1 and its GFP fluorescence are filtered out in this system). SNARF1-loaded BSC emits fluorescence from the cytosol. The brightest pixels areas at 640 nm were selected using FIJI on the basis of their highest fluorescence from a ring closest to the cell contour (presumed to represent the cytosol) and were delimited automatically by yellow lines. The straight lines-delimited area (selected manually) served for background determination. The selected areas were used also for pixel selection in the corresponding 585 nm images and were superimposed for orientation also on the transmitted-light image (Trans). The scale bar: 10 µm. **B.** An *in-situ* calibration of the cytosolic pH in a SNARF1-loaded BSC protoplasts in the presence of the $H^+/K^+$ exchanger, nigericin (5 µM), and a very high $K^+$ conc. in the bath ($[K^+]_{EXT}$ = 200 mM, presumed to equal the cytosolic $[K^+]_{CYT}$; Materials and methods); fluorescence ratio of emitted fluorescence at the indicated wavelengths vs the pH of the buffered bath solution ($pH_{EXT}$). The symbols stand for data from individual cells. The fitted line: F-ratio = 0.32564* pH -1.14968 (Pearson's R = 0.82135); the F-ratio values converted to pH become: pH = (F-ratio +1.14968) / 0.32564.

Suppl. Fig. S4, ABA on BSCs pumps, TT. et al., 2021

Cont. on next page

Cont. from prev. page

**C.** An *in-situ* calibration as in B, after a replacement of the excitation source lamp (and with $[K^+]_{EXT}$ = 250 mM in the bath (Materials and methods); The fitted line: F-ratio= 0.4359*pH -1.9088  (Pearson's R = 0.90397); the F-ratio values can be converted to pH as follows: pH = (F-ratio+1.9088) / 0.4359.

Cont. from prev. page
